# Chemogenetic restoration of autonomous subthalamic nucleus activity ameliorates Parkinsonian motor dysfunction

**DOI:** 10.1101/385443

**Authors:** Eileen L. McIver, Hong-Yuan Chu, Jeremy F. Atherton, Kathleen E. Cosgrove, Jyothisri Kondapalli, David Wokosin, D. James Surmeier, D.Bevan Mark

## Abstract

**Highlights:**

- decorrelating autonomous STN activity was downregulated in both toxin and genetic models of PD
- elevation of D2-striatal projection neuron transmission was sufficient for downregulation
- downregulation was dependent on activation of STN NMDA receptors and _KATP_ channels
- chemogenetic restoration of autonomous spiking reduced synaptic patterning of STN neurons *and* PD motor dysfunction

**eToC:** Excessive synaptic synchronization of STN activity is linked to the symptomatic expression of PD.McIver and colleagues describe the cellular and circuit mechanisms responsible for the loss of decorrelating autonomous STN activity in PD models and demonstrate that chemogenetic rescue of autonomous spiking reduces synaptically patterned STN activity *and* ameliorates Parkinsonian motor dysfunction.

**SUMMARY:** Excessive, synaptically-driven synchronization of subthalamic nucleus (STN) neurons is widely thought to contribute to akinesia, bradykinesia, and rigidity in Parkinson’s disease (PD). Electrophysiological, optogenetic, chemogenetic, genetic, 2-photon imaging, and pharmacological approaches revealed that the autonomous activity of STN neurons, which opposes synaptic synchronization, was downregulated in both toxin and genetic mouse models of PD.Loss of autonomous spiking was due to increased transmission of D2-striatal projection neurons, leading in the STN to elevated activation of NMDA receptors and generation of reactive oxygen species that promoted _KATP_ channel opening.Chemogenetic restoration of autonomous firing in STN neurons reduced synaptic patterning and ameliorated Parkinsonian motor dysfunction, arguing that elevating intrinsic STN activity is an effective therapeutic intervention in PD.

## INTRODUCTION

The glutamatergic subthalamic nucleus (STN) is a small but key component of the basal ganglia, a group of subcortical brain nuclei critical for voluntary movement and a major site of pathology in movement disorders, including Parkinson’s disease (PD) and Huntington’s disease (Albin et al., 1989). Specifically, the STN is a node of the so-called hyperdirect and indirect pathways, which contribute to the initiation, execution, and termination of action sequences through their elevation of GABAergic basal ganglia output (Kravitz et al., 2010; Nambu et al., 2002; Tecuapetla et al., 2016). In PD, increases in the frequency, correlation, and coherence of STN activity have been linked to the expression of akinesia, bradykinesia, and rigidity (Jenkinson and Brown, 2011; Sanders et al., 2013; Sharott et al., 2014; Shimamoto et al., 2013; Zaidel et al., 2009). However, motor symptoms are ameliorated by both inhibition of STN activity through lesions (Bergman et al., 1990) or pharmacological/optogenetic inhibition (Levy et al., 2001; Yoon et al., 2014), and elevation of STN activity through direct electrical stimulation (Benabid et al., 2009; Wichmann et al., 2018), arguing that the abnormal pattern rather than frequency of STN activity impairs movement.Indeed, therapeutic dopamine receptor activation or STN deep brain stimulation decorrelate STN activity/output but do not reverse its hyperactivity (Eusebio et al., 2012; Hashimoto et al., 2003; Wichmann et al., 2018).

In toxin models of PD, abnormal STN activity similar to that seen in patients takes days to weeks to manifest, suggesting that plasticity triggered by the loss of dopamine contributes to its emergence (Mallet et al., 2008b; Vila et al., 2000). Indeed, recent studies in toxin models showed that the strength of external globus pallidus (GPe) inhibition relative to cortical excitation of the STN quadruples through proliferation of GPe synapses (Chu et al., 2015) and removal of cortical synapses (Chu et al., 2017; Mathai et al., 2015). In parallel, the intrinsic autonomous firing of STN neurons is strongly downregulated (Wilson et al., 2006; Zhu et al., 2002a). The available evidence suggests that increased inhibition of the GPe by D2-striatal projection neurons (D2-SPNs) leads to disinhibition of the STN and excessive activation of STN NMDARs, triggering synaptic plasticity (Chu et al., 2015; Chu et al., 2017). Although the cause of autonomous firing downregulation in PD models is unknown, it may be triggered by a similar mechanism because excessive activation of NMDARs can produce oxidant stress and activation of _KATP_ channels, which hyperpolarize STN neurons and inhibit their autonomous spiking (Atherton et al., 2016). Autonomous activity is a fundamental property of the extrastriatal basal ganglia that may reflect the use of GABA by SPNs for encoding and the requirement for tonic inhibitory basal ganglia output to suppress movement (Atherton and Bevan, 2005; Bevan and Wilson, 1999; Chan et al., 2004). In addition, autonomous spiking can decorrelate neuronal activity because the impact of synaptic input depends in large part on the frequency and phase of the postsynaptic oscillatory firing cycle (Baufreton et al., 2005; Bevan et al., 2002; Wilson, 2013). Thus, the loss of autonomous STN activity together with plastic alterations in the strength of synaptic inputs may promote pathological synchronization of the STN and associated motor dysfunction in PD.Using the unilateral 6-hydroxydopamine (6-oHDA)-lesion and genetic Mito-Park models of PD, we addressed 3 hypotheses 1) downregulation of autonomous STN activity is triggered by elevated D2-SPN transmission, leading to disinhibition of the STN and increased activation of STN NMDARs 2) excessive activation of STN NMDARs increases activation of _KATP_ channels, which diminish autonomous activity 3) chemogenetic restoration of autonomous activity reduces the synaptic patterning of STN neurons and ameliorates motor dysfunction.

## RESULTS

Data are reported as median and interquartile range.Unpaired data are represented graphically as box plots, with the median, inter-quartile range, and 10-90% range denoted. Paired data are represented as tilted line segment plots. To minimize assumptions about the distribution of the data, non-parametric, two-tailed statistical tests were applied: Mann-Whitney U (MWU) test for unpaired data, Wilcoxon signed rank (WSR) test for paired comparisons, and Fisher’s exact test for contingency analyses. Exact p-values were corrected for multiple comparisons (Holm, 1979) (**Tables S1-8**).

### Autonomous STN activity is downregulated in PD models

Previous studies have demonstrated that the autonomous firing of STN neurons is disrupted in the 6-hydroxydopamine (6-OHDA) and MPTP toxin models of PD (Chu et al., 2017; Wilson et al., 2006; Zhu et al., 2002a). Furthermore, STN neurons identified as receiving input from the primary motor cortex also exhibited decreased autonomous activity (Chu et al., 2017). We first confirmed the result of Chu et al., 2017 using an identical approach, but a more extensive sample. Thus, in cell-attached patch clamp recordings of STN neurons, the frequency, regularity, and incidence of autonomous firing were lower in *ex vivo* brain slices from unilateral 6-OHDA-injected, dopamine-depleted mice (**Figure 1A-D, H; Table S1**). To determine whether autonomous STN activity is also downregulated in a progressive genetic model of PD, we compared the autonomous spiking of STN neurons in brain slices prepared from 20-week old Mito-Park and littermate control mice.Mito-Park mice are generated by crossing mitochondrial transcription factor A (Tfam)^lox/lox^ (*B6.Cg-Tfam^tm1.1Ncdl^/J*) mice with dopamine transporter (DAT)^cre+/-^ (*B6.SJL-Slc6a3^tm1.1(cre)Bkmn^/J*) mice, which leads to the genetic excision of Tfam in DAT-expressing neurons, the progressive loss of substantia nigra dopaminergic (SN DA) neurons, and development of levo-dopa-sensitive motor dysfunction (Ekstrand and Galter, 2009; Ekstrand et al., 2007). Littermate Tfam^lox/lox^, DAT^cre-/-^ mice were used as controls.At 20 weeks of age, when striatal dopamine has fallen by >95% and the number of tyrosine-hydroxylase expressing SN DA neurons has fallen by >60 *%* (Ekstrand and Galter, 2009; Ekstrand et al., 2007), the frequency, regularity, and incidence of autonomous STN activity in slices from Mito-Park mice were significantly lower than in tissue from control mice (**Figure 1E-H, Table S1**). Together, these data demonstrate that in both acute toxin and progressive genetic models of PD, loss of dopaminergic neuromodulation is associated with the robust downregulation of autonomous STN activity.

**Figure 1.**
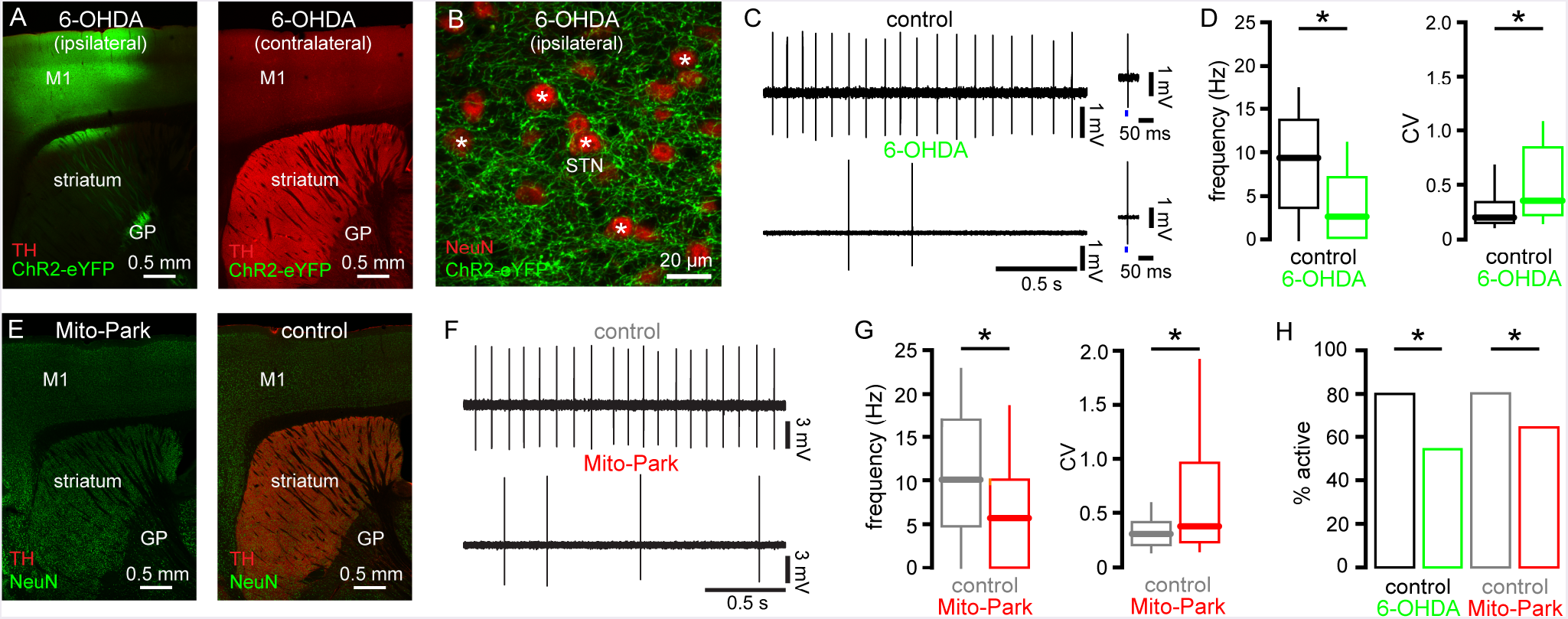
The autonomous activity of STN neurons was downregulated in toxin and genetic models of PD. (A) TyROSine hydroxylase (TH) immunoreactivity (red) and expression of ChR2(H134R)-eYFP (green) ipsilateral and contralateral to injections of 6-OHDA in the MFB and an AAV vector expressing ChR2(H134R)-eYFP in the primary motor cortex (M1). (B) ChR2(H134R)-eYFP (green) expressing motor cortico-STN axon terminals in the vicinity of NeuN-immunoreactive (red) neurons (asterisks) in the ipsilateral STN following the injections described above. (C, D) The frequency and regularity of autonomous activity in STN neurons that receive motor cortical input was reduced in slices from dopamine-depleted mice compared to dopamine-intact control mice (C, examples traces; C insets, response to optogenetic stimulation (blue) of motor cortical input; D, population data). (E) TH immunoreactivity (red) in 20 week-old Mito-Park and control mice (green, NeuN-immunoreactive neurons). (F, G) The frequency and regularity of autonomous activity in STN neurons in slices from Mito-Park mice were reduced relative to age-matched control mice (F, examples traces; G population data). (H) The incidence of autonomous STN activity was significantly reduced in 6-OHDA-injected and Mito-Park mice relative to dopamine-intact control mice.* p < 0.05.ns. See also **Table S1.**

### Elevation of D2-SPN transmission is sufficient to downregulate autonomous STN activity

In PD and its experimental models, loss of striatal DA innervation disinhibits D2-SPNs, which is thought to increase inhibition of GPe and disinhibit the STN (Albin et al., 1989; Lemos et al., 2016; Mallet et al., 2006; Parker et al., 2018; Ryan et al., 2018; Sharott et al., 2017). In mouse toxin models of PD this change in circuit activity is associated with plastic alterations in the relative strength of excitatory and inhibitory synaptic inputs to STN neurons (Chu et al., 2015; Chu et al., 2017; Fan et al., 2012). We therefore hypothesized that downregulation of autonomous STN firing is triggered by Parkinsonian circuit activity.However, SN DA neurons also directly and positively modulate the autonomous activity of STN neurons (Cragg et al., 2004; Loucif et al., 2008; Ramanathan et al., 2008; Zhu et al., 2002a; Zhu et al., 2002b), suggesting that reduced STN activity in PD models could result from the loss of local DA signaling. To determine whether an increase in indirect pathway activity is sufficient to downregulate autonomous STN activity, D2-SPNs were chemogenetically stimulated in dopamine-intact BAC transgenic B6.Cg-Tg(Adora2a-Chrm3*,-mCherry)AD6Blr/J (adora2A-rM3D(Gs)-mCherry) mice that express rM3D(Gs)-mCherry in D2-SPNs under the control of the A2A-promoter, as described previously (Bouabid and Zhou, 2018; Chu et al., 2017; Farrell et al., 2013). Consistent with the chemogenetic activation of D2-SPNs in these mice 1) subcutaneous (SC) administration of 1 mg/kg clozapine-N-oxide (CNO) reduces motor activity in the open field and the frequency of GPe activity *in vivo* (Bouabid and Zhou, 2018; Chu et al., 2017; Farrell et al., 2013) 2) application of 10 μM CNO to brain slices elevates the frequency of GABAergic synaptic currents in GPe neurons (Chu et al., 2017). The frequency, regularity, and incidence of autonomous STN activity in *ex vivo* brain slices from CNO-treated adora2A-rM3D(Gs)-mCherry mice (1 mg/kg every 12 hours for 3 days) was significantly reduced compared to activity in slices from vehicle-treated mice (**Figure 2A-D, Table S2**). In regular C57BL/6 mice that lack expression of rM3D(Gs)-mCherry, chronic CNO administration had no effect on autonomous activity compared to activity in CNO-naïve dopamine-intact mice (**Figures 1A-D, 1H and S1A-B; Tables S1 and S2**). Together these data demonstrate that elevation of D2-SPN transmission is sufficient to trigger the downregulation of autonomous STN activity.

**Figure 2.**
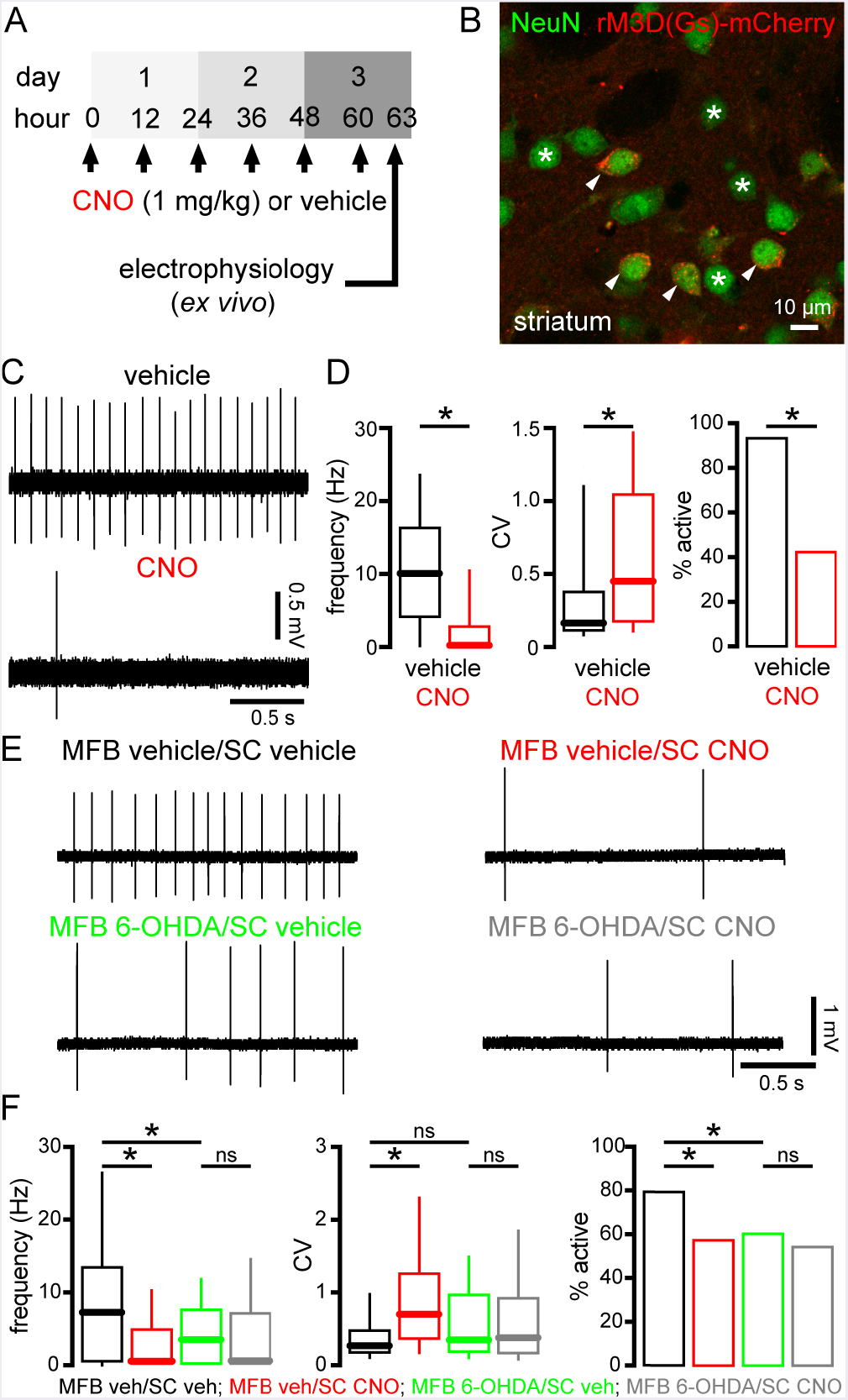
Chemogenetic activation of D2-SPNs was sufficient to downregulate autonomous STN activity in dopamine-intact mice but this effect was occluded in dopamine-depleted mice. (A) Schedule of SC CNO or vehicle injection and *ex vivo* electrophysiological recording in adora2a-rM3D(Gs)-mCherry mice. (B) Expression of rM3D(Gs)-mCherry (red) in a subset of NeuN-immunoreactive striatal neurons (green) in an adora2a-rM3D(Gs)-mCherry mouse (arrows: expressing; asterisks: non-expressing).(C, D) The autonomous activity of STN neurons was downregulated in slices from CNO-compared to vehicle-treated adora2a-rM3D(Gs)-mCherry mice (C, example traces; D, population data). (E, F) In MFB vehicle-injected adora2a-rM3D(Gs)-mCherry mice, autonomous STN activity was downregulated in slices from CNO-compared to vehicle-treated mice. However, in MFB 6-OHDA-injected adora2a-rM3D(Gs)-mCherry mice, autonomous STN activity was downregulated similarly in slices from CNO- and vehicle-treated mice, arguing that the effect of chemogenetic stimulation of D2-SPNs was occluded by dopamine depletion. (E, example traces; F, population data). * p < 0.05.ns, not significant. See also **Table S2and Figure S1**

If elevated striato-pallidal transmission and subsequent disinhibition of the STN underlies downregulation of autonomous STN activity in 6-OHDA-treated mice, the effect of chemogenetic activation of D2-SPNs in adora2ArM3D(Gs)-mCherry mice should be occluded by dopamine depletion. To test this hypothesis, adora2A-rM3D(Gs)-mCherry mice were injected unilaterally in the MFB with either 6-OHDA or vehicle. Approximately 2-3 weeks later, mice received SC injections of either vehicle or CNO and the autonomous activity of STN neurons from the hemisphere ipsilateral to the MFB injection was analyzed, as described above. Consistent with our hypothesis 1) autonomous firing was downregulated by 6-OHDA injection, as evinced by comparison of data from SC vehicle-treated mice with vehicle or 6-OHDA MFB injection 2) autonomous firing was downregulated following chemogenetic activation of D2-SPNs in dopamine intact mice, as evinced by comparison of data from MFB vehicle-injected mice with SC vehicle or CNO treatment 3) the downregulatory effect of chemogenetic stimulation of D2-SPNs on autonomous STN activity was occluded by dopamine depletion, as evinced by the similarity of data from MFB 6-OHDA-injected mice with SC vehicle or CNO treatment (**Figure 2E, F; Table S2**). Together the data are consistent with the conclusion that elevated D2-SPN transmission contributes to the downregulation of autonomous STN activity in 6-OHDA-injected mice.

### NMDARs are necessary for the downregulation of autonomous STN activity in PD mice

In 6-OHDA-treated mice, disinhibition of the STN due to elevated D2-SPN transmission leads to NMDAR-dependent alterations in the relative number and strength of inhibitory and excitatory synaptic inputs (Chu et al., 2015; Chu et al., 2017). Therefore, increased activation of STN NMDARs in 6-OHDA-injected mice might also be responsible for the downregulation of autonomous STN activity.Indeed, pre-incubation of brain slices from dopamine-intact mice with 25 μM NMDA for 1 hour was sufficient to persistently downregulate autonomous STN activity (Atherton et al., 2016). To test the hypothesis that downregulation of autonomous STN activity in 6-OHDA-injected mice is dependent on NMDARs, STN NMDARs were knocked down by stereotaxic injection of an adeno-associated virus (AAV) carrying a cre-recombinase expression construct into the STN of mice in which *GRIN1*, the gene encoding the obligatory subunit of the NMDAR, was floxed at both alleles (Tsien et al., 1996) (B6.129S4-*Grin1^tm2Stl^/J*; **Figure 3A-D, Table S3**). To control for viral infection another set of *GRIN1^lox/lox^* mice received injection of an AAV carrying only an eGFP reporter construct (**Figure 3A-D, Table S3**). Viral injections were performed at the time as 6-OHDA or vehicle injection in the MFB. Consistent with our hypothesis, knockdown (KD) of STN NMDARs prevented the downregulation of autonomous STN activity in 6-OHDA-injected mice (**Figure 3C-D, Table S3**). If activation of STN NMDARs *in vivo* is responsible for the downregulation of autonomous STN activity in 6-OHDA-injected mice, then the downregulatory effect of NMDAR activation *ex vivo* on autonomous STN activity should be occluded in slices from 6-OHDA-injected but not vehicle-injected C57BL/6 mice. Consistent with this hypothesis, we found that 1) the autonomous firing of STN neurons in untreated slices from 6-OHDA-injected mice was downregulated relative to that in untreated slices from vehicle-injected mice (**Figure 3E-F, Table S3**) 2) NMDAR activation *ex vivo* downregulated autonomous STN activity in slices from vehicle-but not 6-OHDA-injected mice relative to untreated slices (**Figure 3E-F**). Together these data suggest that STN NMDARs are necessary for the downregulation of autonomous STN activity in 6-OHDA-injected mice.

**Figure 3.**
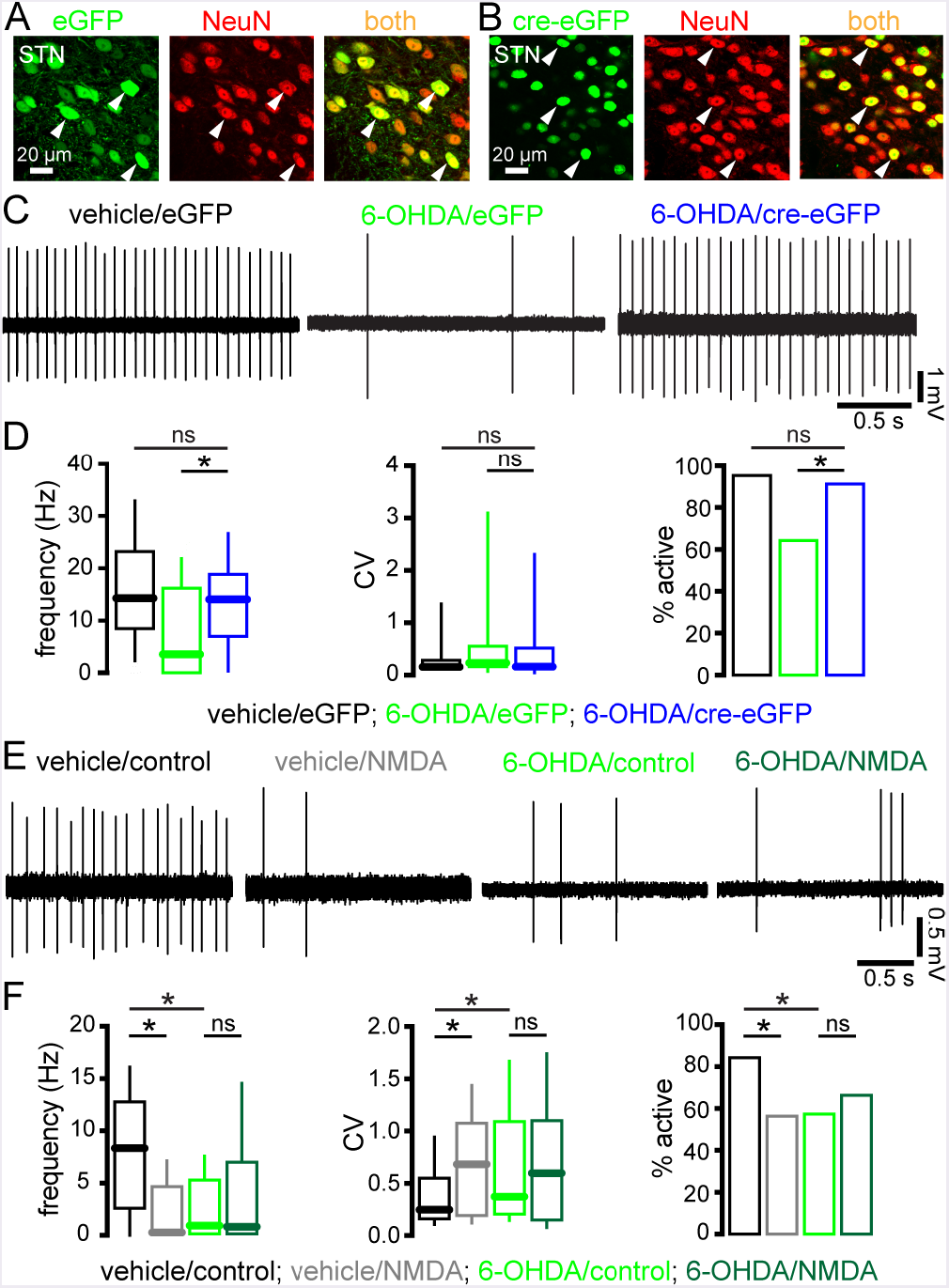
Downregulation of autonomous STN activity in dopamine-depleted mice was dependent on activation of STN NMDARs *in vivo*. (A,B) Confocal micrographs of viral-mediated eGFP green) or cre-eGFP (B, green) expression in NeuN-immunoreactive (red) STN neurons (arrowheads) in *Grin1^lox/lox;^* mice.(C, D) In eGFP-expressing, NMDAR-intact STN neurons autonomous activity was downregulated in brain slices from 6-OHDA-injected mice relative to activity in vehicle-injected mice. However, in cre-eGFP-expressing, NMDAR KD STN neurons downregulation of autonomous activity in 6-OHDA-injected mice was prevented (C, examples; D, population data). (E, F) Consistent with **Figures 1** and **2**, the autonomous firing of STN neurons in untreated slices from 6-OHDA-injected, NMDAR-intact mice was downregulated relative to that in untreated slices from vehicle-injected, NMDAR-intact mice.Activation of STN NMDARs *ex vivo* downregulated autonomous activity in slices from vehicle-but not 6-OHDA-injected, NMDAR-intact mice, arguing that in 6-OHDA-injected mice NMDAR activation *in vivo* occluded the downregulatory effect of NMDAR activation *ex vivo* (E, examples; F, population data). * p < 0.05.ns, not significant. See also **Table S3**

### STN neurons exhibit elevated oxidant stress and hydrogen peroxide-triggered K_ATP_ channel activation in PD mice

In STN neurons, NMDAR (and mGluR) activation can stimulate the opening of ATP-sensitive K+ (K_ATP_) channels, which limit activity (Atherton et al., 2016; Shen and Johnson, 2010, 2013). NMDAR activation can promote Katp channel opening through several mechanisms 1) a fall in the ratio of cytosolic ATP to ADP (Nichols, 2006; Wang and Michaelis, 2010) 2) the direct actions of reactive oxygen and/or nitrogen species (RoS and RNS, respectively) (Atherton et al., 2016; Ichinari et al., 1996; Kawano et al., 2009; Lee et al., 2015) 3) neuromodulation by ROS- and RNS-linked second messenger pathways (Shen and Johnson, 2010; Zhang et al., 2014). Indeed, we showed recently that chronic activation of STN NMDARs can persistently downregulate autonomous activity through ROS-dependent K_ATP_ channel opening in HD mice, due to impaired astrocytic glutamate uptake, and in WT mice, through application of exogenous NMDA *ex vivo* (Atherton et al., 2016). To determine whether the NMDAR-dependent downregulation of autonomous STN activity in 6-OHDA-injected mice is dependent on ROS-dependent activation of K_ATP_ channels, we conducted a series of experiments. To determine whether disinhibition of STN neurons in 6-OHDA-injected mice elevates oxidant stress, we virally expressed the mitochondrial matrix-targeted thiol redox sensor mito-roGFP in STN neurons (Atherton et al., 2016; Hanson et al., 2004). Calibrated measurements of mito-roGFP fluorescence using 2-photon laser scanning micROScopy (2-PLSM) confirmed that oxidant stress in STN mitochondria was elevated in slices from 6-OHDA-injected mice (**Figure 4A, B, Table S4**) compared with those from vehicle-injected controls (**Figure 4A, B, Table S4**). To determine whether the elevation of ROS could cause the downregulation of autonomous STN activity, a membrane permeable variant of catalase, which metabolizes hydrogen peroxide, was applied to *ex vivo* brain slices.Catalase application rapidly reversed the deficit in STN activity in brain slices from 6-OHDA-injected mice (**Figure 4C, D, Table S4**). Although this treatment also modestly increased the frequency and regularity of firing in slices from vehicle-injected mice, the effects were significantly smaller than in 6-OHDA-injected mice.To determine whether K_ATP_ channels contribute to the downregulation of autonomous STN firing, we next compared the effect of the K_ATP_ channel inhibitor glibenclamide (100 nM) on the autonomous activity of STN neurons in slices from 6-OHDA- and vehicle-injected mice.Glibenclamide rescued the frequency and regularity of autonomous STN activity in brain slices from 6-OHDA-injected mice; although this treatment also modestly increased the frequency and regularity of firing in slices from vehicle-injected mice, the effects were significantly greater in 6-OHDA-injected mice (**Figure 4E, F, Table S4**). The effects of catalase or K_ATP_ channel inhibition on spiking were similar, suggesting that they were mediated through a common pathway (**Figure 4G, Table S4**). Consistent with this inference, the effects of catalase on STN spiking were occluded by prior K_ATP_ channel inhibition with glibenclamide (**Figure 4H, I Table S4**). To determine whether ATP depletion was also a factor, the spiking rate in cell-attached recordings was compared to that after breaking into the cell with an internal solution containing 2 mM ATP.Dialysis with ATP did not change spiking in STN neurons from lesioned mice, arguing that ATP depletion was not a factor in downregulating autonomous activity (**Figure 4J, K, Table S4**).

**Figure 4.**
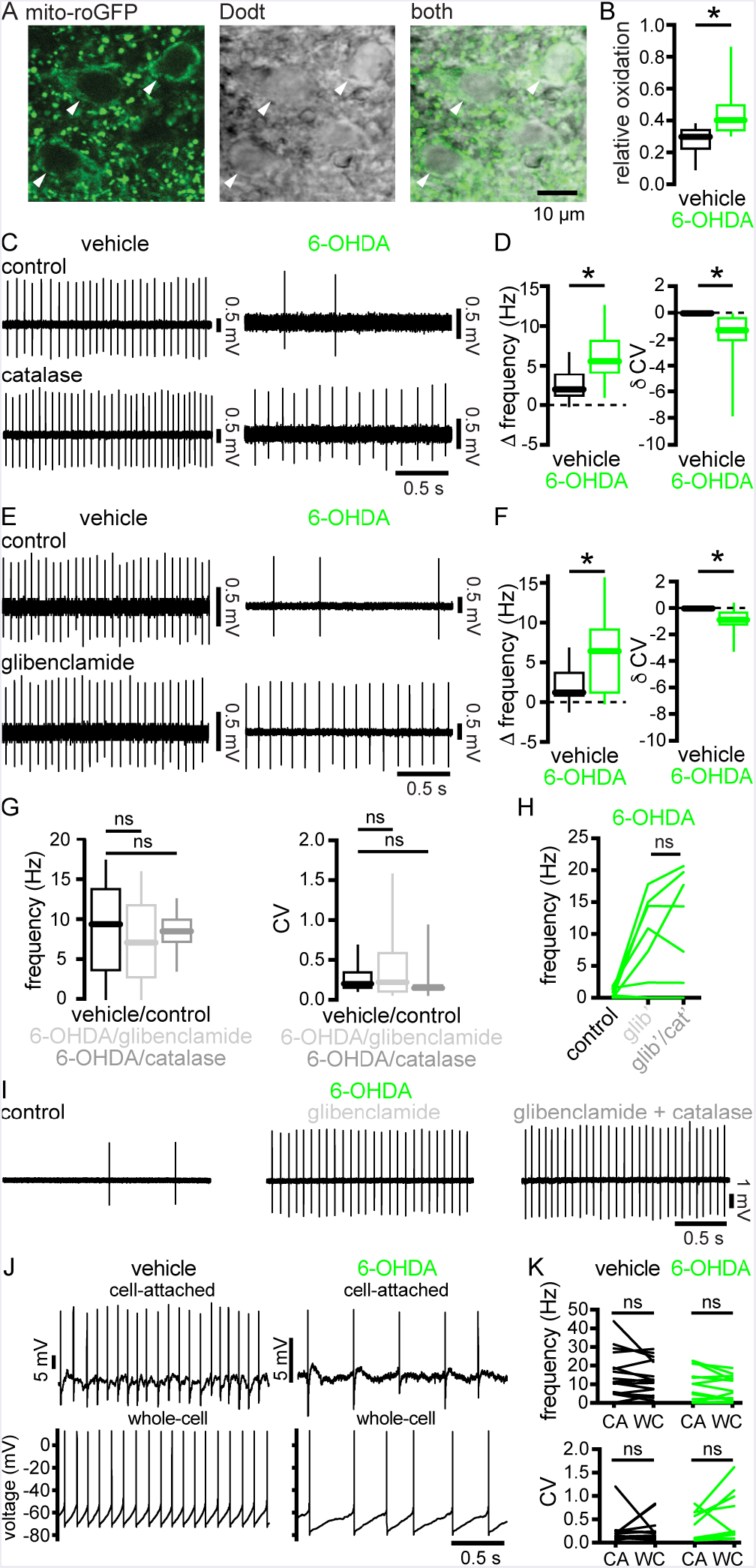
STN neurons exhibited elevated mitochondrial oxidation and hydrogen peroxide-triggered _K_ATP__ channel activation in PD mice. (A) The expression of mito-roGFP (green) in STN neurons (arrowheads) and their somatic morphology (grayscale) were imaged under simultaneous 2-photon laser scanning fluorescent and Dodt contrast micROScopy, respectively. (B) The relative oxidation of mitochondria in STN neurons was elevated in slices from 6-OHDA-versus vehicle-injected mice. (C, D) Breakdown of hydrogen peroxide with catalase rescued autonomous STN activity in slices from 6-OHDA-injected mice but had relatively minimal effects on neurons from vehicle-injected mice (C, examples; D, population data). (E, F) Inhibition of K_ATP_ channels with glibenclamide rescued autonomous STN activity in slices from 6-OHDA-injected mice but had relatively minimal effects on neurons from vehicle-injected mice (E, examples; F, population data). (G) Autonomous firing in slices from 6-OHDA-injected mice that had been treated with glibenclamide or catalase were not significantly different from autonomous firing in slices from dopamine-intact control mice (**Figure 1C, D**). (H, I) The effect of catalase on autonomous STN firing in slices from 6-OHDA-injected mice was occluded by prior application of glibenclamide (H, population data; I, example). (J, K) In slices from vehicle- or 6-OHDA-injected mice autonomous STN activity initially recorded in the cell-attached configuration was not altered by subsequent establishment of the whole-cell configuration with patch pipettes containing 2 mM ATP (J, example; K, population data). * p < 0.05.ns, not significant. See also **Figures S2** and **S3** and **Table S4**

We next tested whether the downregulation of autonomous STN firing produced by chronic chemogenetic stimulation of D2-SPNs was also mediated through K_ATP_ channels. Consistent with this mechanism, glibenclamide rescued the frequency and regularity of autonomous STN firing in slices from CNO-treated mice, but had minimal effects in vehicle-treated mice (**Figure S2A, B, Table S4**). Finally, to determine whether the downregulation of autonomous STN firing in Mito-Park mice was also mediated through K_ATP_ channels, we compared the effect of glibenclamide on autonomous STN activity in 20-week Mito-Park and litter-mate control mice. Inhibition of K_ATP_ channels rescued autonomous spiking in slices from Mito-Park mice, while having relatively minimal effects in slices from control mice (**Figure S2C, D, Table S4**). Taken together, the data argue that following the loss of dopamine, disinhibition of the STN leads to increased activation of STN NMDARs, which increases ROS generation and K_ATP_ channel opening, resulting in the downregulation of autonomous STN activity.

It was recently reported that the expression of small-conductance Ca^2+^-activated K_+_ (SK) channels, which are important regulators of the pace and precision of autonomous STN activity (Hallworth et al., 2003), is upregulated in 6-OHDA-injected rodents (Mourre et al., 2017). To determine their role, the effect of 10 nM apamin (an SK channel blocker) was examined. Inhibition of SK channels did not rescue autonomous STN activity in slices from 6-OHDA-treated mice (**Figure S3**). Furthermore, the effects of SK channel blockade on autonomous STN activity were similar in 6-OHDA- and vehicle-injected mice, arguing that SK channels were not responsible for the downregulation of autonomous activity in 6-OHDA-injected mice.

### Chemogenetic restoration of autonomous STN activity reduces cortico-STN patterning

To assess how the loss of autonomous STN firing affects synaptic patterning and motor dysfunction in PD mice, a chemogenetic strategy (Roth, 2016) was used to restore intrinsic activity (**Figures 5-7, Tables S5-S7**). An AAV vector carrying a synapsin promoter-driven hM3D(Gq)-mCherry construct was injected into the STN ipsilateral to an injection of 6-OHDA or vehicle in the MFB (**Figure 5A**). 2-3 weeks later brain slices were prepared and the effect of chemogenetic activation of hM3D(Gq)-mCherry with 10 μM CNO on intrinsic STN activity was tested. CNO application increased the frequency and regularity of autonomous STN activity in slices from both 6-OHDA- and vehicle-injected mice, presumably due to the generation of a tonic inward current that could be measured under voltage clamp at -60 mV (**Figure 5B-E, Table S5**). These effects occurred within a minute of CNO application and persisted for the remainder of the recording, which was usually a further 10-30 minutes. No effect on the amplitude or short-term plasticity of electrically evoked postsynaptic currents were observed, indicating that chemogenetic activation did not directly affect afferent synaptic transmission (**Figure 5F, G, Table S5**). These data demonstrate that chemogenetic activation of hM3D(Gq)-mCherry promoted intrinsic, autonomous STN activity.

**Figure 5.**
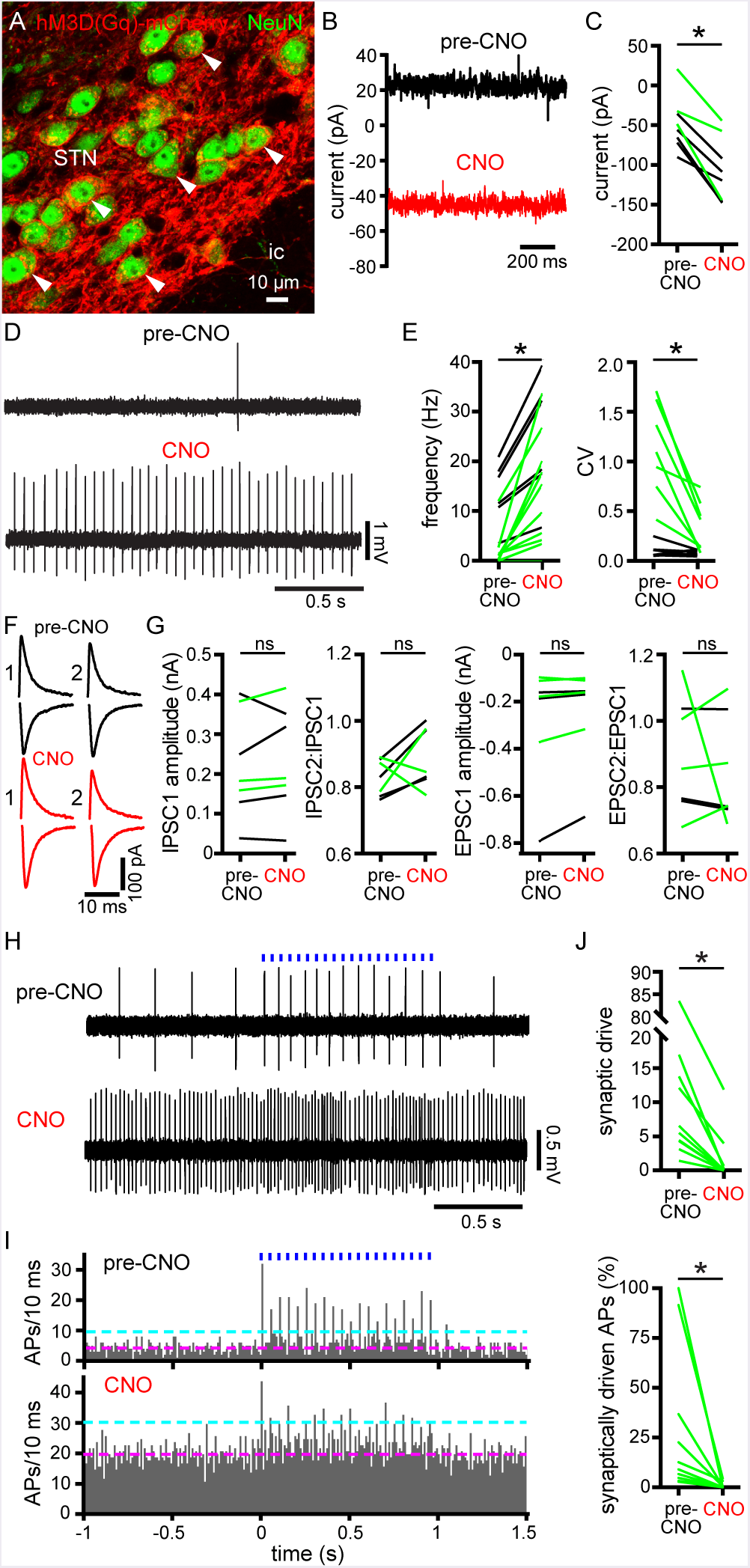
(A) Chemogenetic rescue of autonomous firing reduced synaptic patterning of STN neurons in brain slices from 6-OHDA-injected mice. (A)Confocal micrograph of expression of hM3D(Gq)-mCherry (red) in NeuN-immunoreactive STN neurons.(B, C) Chemogenetic activation of hM3D(Gq)-mCherry with 10 μM CNO generated a tonic inward current that could be measured under voltage clamp at -60 mV in STN neurons from 6-OHDA- (green) and vehicle- (black) injected mice (B, example from a 6-OHDA-injected mouse; C, population). (D, E) Chemogenetic activation of hM3D(Gq)-mCherry increased the frequency and regularity of autonomous STN activity in *e× viv¤* brain slices from 6-OHDA- (green) and vehicle- (black) injected mice (D, example from a 6-OHDA-injected mouse; E, population). (F, G) Pairs of I PSCs (upper black traces; holding voltage, -60 mV) and EPSCs (lower red traces; holding voltage, -80 mV) electrically evoked in STN neurons with an interval of 100 ms were unaffected by chemogenetic activation of hM3D(Gq)-mCherry (F, examples from a 6-OHDA-injected mouse; G, population data; black/green data from vehicle-/6-OHDA-injected mice, respectively). (H-J) The patterning of STN activity by motor cortical inputs optogenetically stimulated at 20 Hz for 1 second (blue) in *ex vivo* brain slices from 6-OHDA-injected mice was reduced by chemogenetic activation of hM3D(Gq)-mCherry in STN neurons (H, example cell-attached recordings; I, population peristimulus time histogram (magenta line, mean; cyan line, mean + 3SD); J, population data; data was derived from 5 trials of stimulation per neuron).*, p < 0.05.ns, not significant. See also **Table S5**

Next, the effect of restoring the autonomous activity of STN neurons on the response to cortical stimulation was examined. Abnormally correlated, often coherent activity in the cortex and STN in PD and its experimental models has been consistently linked to motor dysfunction (Mallet et al., 2008b; Sharott et al., 2014; Shimamoto et al., 2013; Zaidel et al., 2009). Autonomous firing of STN neurons 1) opposes synchronization of STN activity because the effect of synaptic input is dependent on its timing relative to the postsynaptic neuron’s firing cycle 2) generates a basal level of synaptic output from the STN that synaptic input can add to and/or subtract from and/or adjust the timing of (Bevan et al., 2002; Wilson, 2013). We therefore hypothesized that chemogenetic rescue of autonomous STN activity in slices from 6-OHDA-injected mice should diminish synaptic patterning of STN activity evoked by optogenetic stimulation of motor cortical inputs. ChR2(H134R)-eYFP was expressed virally in motor cortical projections to the STN through AAV injection in the motor cortex, as described previously (Chu et al., 2015; Chu et al., 2017).*Ex vivo* brain slices were prepared 2-3 weeks later and cortical inputs were stimulated using 1 ms duration blue light pulses delivered at 20 Hz for 1 second (**Figure 5H, I**). The number of “synaptically-driven” action potentials during optogenetic stimulation of cortico-STN inputs was defined as the number of spikes that exceeded the mean + 3SD of action potentials in the second prior to stimulation (**Figure 5J**). Consistent with our hypothesis, chemogenetically generated intrinsic STN activity profoundly reduced the number of cortically driven spikes both in absolute terms and as a percentage of those generated intrinsically in each neuron tested (**Figure 5H-J**).

### Chemogenetic restoration of autonomous STN activity ameliorates motor dysfunction in dopamine-depleted mice

Interruption of abnormal STN activity through lesioning, pharmacological inactivation, or deep-brain electrical stimulation ameliorates motor dysfunction in PD and its models (Benabid et al., 2009; Bergman et al., 1990; Levy et al., 2001; Wichmann et al., 2018; Yoon et al., 2014). The fact that both decreasing and increasing STN activity are therapeutic argues that disrupting the aberrant pattern of STN activity, rather than lowering its mean rate, is key to symptomatic relief. Furthermore, STN NMDAR KD, which prevents the loss of autonomous STN activity (and the decrease in strength of synaptic excitation relative to inhibition of STN neurons) following dopamine depletion, ameliorates motor dysfunction (Chu et al., 2017). We therefore hypothesized that chemogenetic restoration of autonomous activity would improve motor function in 6-OHDA-injected mice. The impact of chemogenetic activation of virally expressed hM3D(Gq)-mCherry in the STN on the motor function of unilateral 6-OHDA-injected mice was therefore tested. Injection of AAV expressing of hM3D(Gq)-mCherry in the STN and 6-OHDA or vehicle in the MFB were performed, as described above. To confirm that CNO-administration led to the chemogenetic activation of STN neurons and their target structures *in vivo*, CNO (1 mg/kg) or vehicle was administered subcutaneously to MFB-vehicle- and -6-OHDA-injected mice with viral expression of hM3D(Gq)-mCherry in the STN. Ninety minutes later, mice were perfused transcardially with an aldehyde-containing fixative and their brains were processed for immunohistochemical detection of 1) c-fos, an immediate early gene whose expression is associated with elevated firing (Morgan et al., 1987) 2) parvalbumin, a Ca^2^+-binding protein whose expression in the SNr can be used to identify GABAergic SN basal ganglia output neurons (Rajakumar et al., 1994). Consistent with the chemogenetic activation of STN neurons, the density of STN neurons that expressed c-fos was greater in tissue from CNO-treated mice (**Figure 6A-C, Table 6**). Furthermore, the density of PV-immunoreactive SNr neurons that expressed c-fos was greater in tissue from CNO-treated mice, consistent with an increase in synaptic drive from the STN (**Figure 6D-F, Table 6**)

**Figure 6.**
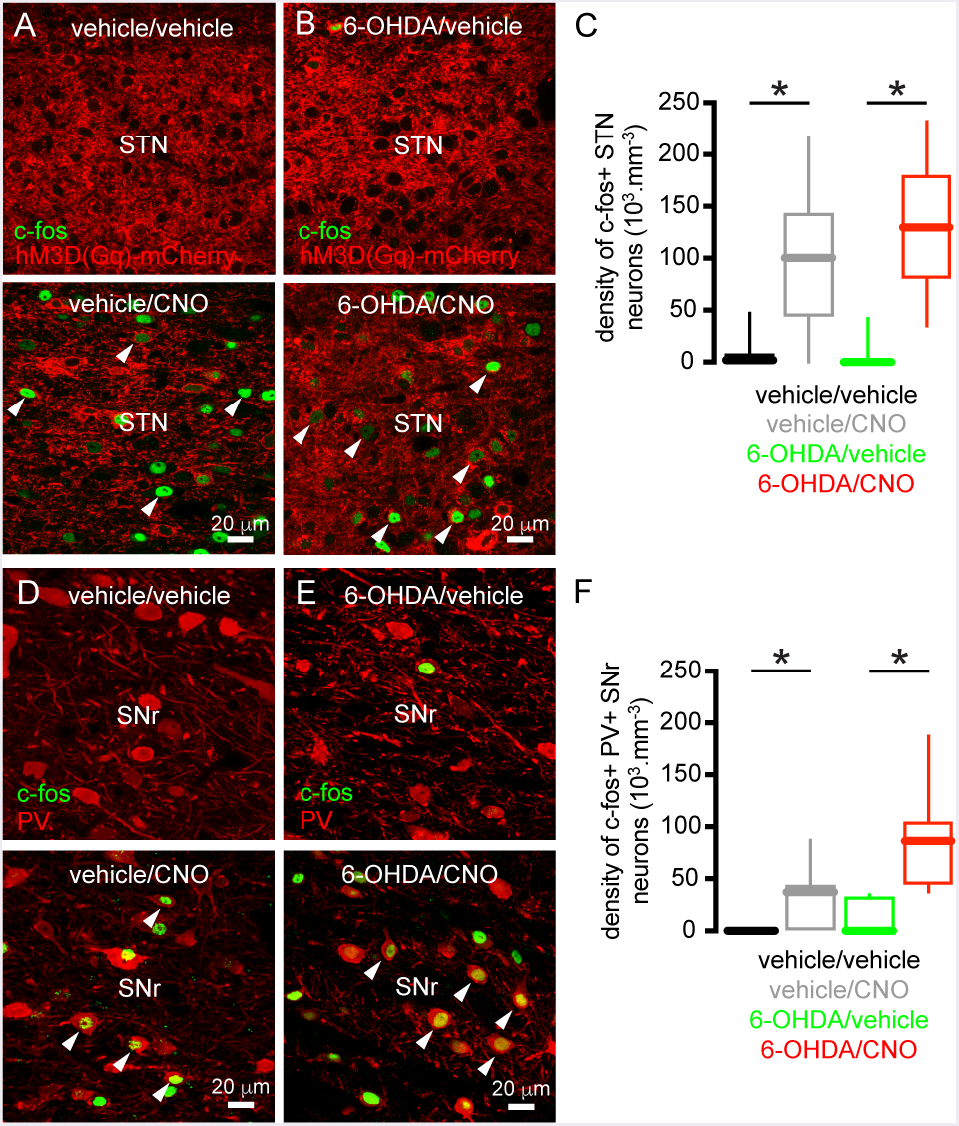
Chemogenetic activation of STN neurons *in vivo* increased immediate early gene expression in the STN and its targets. (A-C) The density of hM3D(Gq)-mCherry-expressing STN neurons that expressed c-fos was elevated in dopamine-intact and -depleted mice that were treated with CNO versus those treated with vehicle (A, B, examples; C, population data).(D-F) The density of PV-immunoreactive SN neurons that expressed c-fos was elevated in dopamine-intact and -depleted mice that were treated with CNO versus those treated with vehicle (D, E, examples; F, population data).* p < 0.05. See also **Table S6**

The impact of chemogenetic activation of the STN on motor function in 6-OHDA-injected dopamine-depleted mice was assessed in a glass cylinder during vertical exploration. When placed in a small-diameter cylinder, rodents rear and in so doing place their forelimbs on the sides of the cylinder (Schallert et al., 2000). In unilateral MFB vehicle-injected, dopamine-intact mice forelimb usage was symmetric and chemogenetic activation of the STN (SC, 1 mg/kg CNO) had no consistent effect (**Fig.7A, B**). In unilateral MFB 6-OHDA-injected, dopamine-depleted rodents, forelimb usage was asymmetric: forelimb use contralateral to the 6-OHDA toxin injection was reduced relative to the ipsilateral forelimb *and* to contralateral forelimb usage in MFB vehicle-injected, dopamine-intact mice, as described previously (Schallert et al., 2000) (**Figure 7A-D; Movie S1**). Within 30 minutes of chemogenetic stimulation of the STN (SC, 1 mg/kg CNO), unilateral 6-OHDA-injected mice with ipsilateral hM3D(Gq)-mCherry expression in the STN increased their relative usage of the contralateral Parkinsonian forelimb *and* decreased their relative usage of the ipsilateral forelimb compared to baseline usage (**Figure 7C; Movie S1**). Twenty-four hours after CNO injection, forelimb usage in unilateral 6-OHDA-injected, dopamine-depleted mice had returned to its baseline asymmetric state (data not shown). In unilateral MFB 6-OHDA-injected, dopamine-depleted mice, with hM3D(Gq)-mCherry expressed ipsilaterally in regions adjacent to but not including the STN, chemogenetic activation also increased contralateral forelimb usage compared to baseline activity (**Figure 7D**). However, the rescue of contralateral forelimb usage following chemogenetic activation of the STN was greater than that produced by chemogenetic activation of structures surrounding the STN (**Figure 7E**). Together these data demonstrate that chemogenetic rescue of autonomous STN activity in unilateral 6-OHDA-injected mice increases usage of the contralateral Parkinsonian forelimb during vertical exploratory behavior.

**Figure 7.**
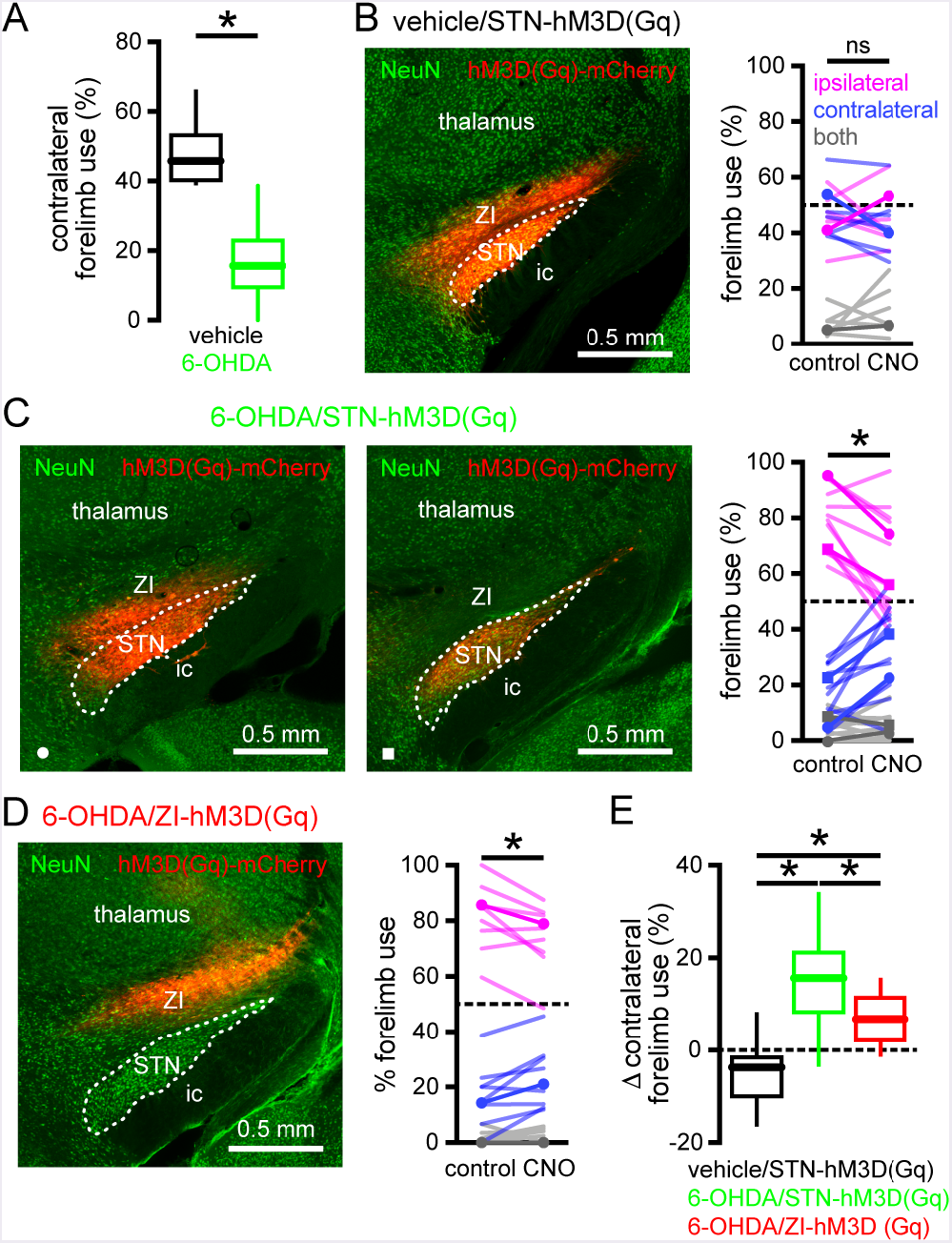
Chemogenetic activation of the STN improved motor function in unilateral 6-OHDA-injected mice. (A) In unilateral 6-OHDA-injected mice (green) contralateral forelimb use during vertical exploration was significantly lower than in dopamine-intact, vehicle-injected mice (black).(B, E) In vehicle-injected, dopamine-intact mice forelimb use was symmetric and not affected by chemogenetic activation (CNO) of hM3D(Gq)-mCherry expressed in the STN and zona incerta (ZI) (left panel, hM3D(Gq)-mCherry expression; right panel population data, example highlighted).(C, E) In 6-OHDA-injected, dopamine-depleted mice forelimb usage was asymmetric but ameliorated by chemogenetic activation of hM3D(Gq)-mCherry expressed in the STN and ZI (left panel) or the STN alone (middle panel; expression was restricted to the STN in a GABRR3-cre mouse using injection of an AAV expressing a cre-dependent hM3D(Gq)-mCherry construct; right panel, population data with examples highlighted).(D, E) In 6-OHDA-injected, dopamine-depleted mice asymmetric forelimb use was moderately ameliorated by chemogenetic activation of hM3D(Gq)-mCherry expressed in the ZI (panels as for B).(E) However, the chemogenetic rescue of forelimb usage in 6-OHDA-injected mice was significantly greater when hM3D(Gq)-mCherry expression involved the STN.* p < 0.05.ns, not significant. See also **Table S7** and **Video S1**.

## DISCUSSION

Autonomous firing is a fundamental property of STN neurons, which is generated by an interplay of ion channels in the axon initial segment and somatodendritic region, and produces a basal level of activity and synaptic output that can be bi-directionally adjusted by incoming synaptic input (Atherton et al., 2008; Bevan and Wilson, 1999; Do and Bean, 2003). Autonomous firing also helps to decorrelate neuronal activity across the STN because the effect of synaptic input is dependent on its timing relative to the phase of postsynaptic oscillatory firing cycle, which is generated independently in each neuron (Baufreton et al., 2005; Bevan et al., 2002; Wilson, 2013). In both the 6-OHDA and Mito-Park models of PD, the autonomous activity of STN neurons was downregulated, which may promote synaptic synchronization and increase the amount of synaptically patterned activity relative to that generated intrinsically. Excessive synaptic synchronization of STN neurons in PD is thought to degrade the information coding capacity of the nucleus and thus limit its capability to transmit complex signals necessary for the execution of action sequences (Mallet et al., 2008b; Wilson, 2013).

In PD models, loss of inhibitory D2 receptor DA neuromodulation leads to an increase in the frequency and synchronization of D2-SPN activity and synaptic output (Lemos et al., 2016; Mallet et al., 2006; Parker et al., 2018; Ryan et al., 2018; Sharott et al., 2017). These changes have been attributed to disinhibition of D2-SPN dendrites and synapses and remodeling of cortical and fast-spiking interneuron inputs (Day et al., 2006; Fieblinger et al., 2014; Gittis et al., 2011; Lemos et al., 2016). Thus, in response to phasic cortical activity, D2-SPNs produce abnormally strong, phasic suppression of GPe activity at the same time that the STN is subjected to direct cortical excitation (Magill et al., 2001; Mallet et al., 2008a; Walters et al., 2007; Zold et al., 2007). In dopamine-intact mice, we found that chemogenetic stimulation of D2-SPNs was sufficient to downregulate autonomous STN activity, arguing that the loss of autonomous activity in PD models may be driven by an increase in D2-SPN activity/transmission. Consistent with this inference, prior 6-OHDA lesioning occluded the downregulatory effect of chemogenetically stimulating D2-SPNs. Although the loss of dopaminergic neuromodulation within the STN (Loucif et al., 2008; Ramanathan et al., 2008; Zhu et al., 2002b) cannot be excluded as a factor, the extent to which STN activity was downregulated in dopamine-intact mice was reproduced solely by D2-SPN stimulation. The proposition that disinhibition drives the downregulation of STN spiking is also consistent with the observation that KD of STN NMDARs prevented the effect of dopamine depletion on autonomous STN activity. Furthermore, the ability of NMDAR stimulation to suppress STN spiking *ex vivo* (Atherton et al., 2016) was occluded in slices from 6-OHDA-lesioned mice, suggesting that this mechanism was already engaged. These data, together with recent studies of synaptic excitation (Chu et al.,2015) and inhibition (Chu et al., 2017), reveal that the level of autonomously generated STN activity *and* the balance of synaptic excitation and inhibition of STN neurons are regulated by STN NMDARs.

How do NMDARs regulate autonomous STN activity? NMDAR activation can stimulate mitochondrial respiration through several mechanisms, including mitochondrial Ca^2^+ loading or a reduction in the cytosolic ATP/ADP ratio, and thus increase their production of ROS (Dugan et al., 1995; Stanika et al., 2010). Indeed, in STN neurons from 6-OHDA-injected mice, mitochondrial matrix oxidant stress was chronically elevated. Elevated mitochondrial oxidation could also impair production of ATP and thus limit autonomous firing through an ATP-dependent mechanism (Wang and Michaelis, 2010). However, dialysis of STN neurons with ATP had no effect. Instead breakdown of the ROS hydrogen peroxide rapidly reversed autonomous firing deficits in slices from dopamine-depleted mice, while having relatively minor effects in slices from dopamine-intact mice. K_ATP_ channels are key metabolic sensors that limit cellular excitability in response to a reduction in ATP to ADP ratio and oxidative stress (Lee et al., 2015; Nichols, 2006). In dopamine-depleted mice, we found that inhibition of K_ATP_ channels restored autonomous STN activity and occluded the effect of hydrogen peroxide breakdown, while having relatively modest effects in tissue from dopamine-intact mice. Because hydrogen peroxide directly promotes K_ATP_ channel opening (Ichinari et al., 1996; Lee et al., 2015) and breakdown of hydrogen peroxide rapidly reversed firing deficits, these data suggest that hydrogen peroxide is directly responsible for increased activation of K_ATP_ channels in STN neurons from PD mice, rather than e.g., RNS-dependent modulation of K_ATP_ channels (Kawano et al., 2009; Shen and Johnson, 2010). Indeed, in dopamine-intact mice elevation of hydrogen peroxide through inhibition of glutathione peroxidase reduces autonomous STN activity through activation of K_ATP_ channels (Atherton et al., 2016). Interestingly, inhibition of K_ATP_ channels also restored autonomous firing in Mito-Park mice and in dopamine-intact mice following chemogenetic stimulation of D2-SPNs, pointing to a common mechanism of downregulation in 3 different models of parkinsonism. Precisely why ROS levels and K_ATP_ channel activation remain persistently elevated in *ex vivo* brain slices prepared from these mouse models is unclear but possible mechanisms include 1) increased ROS production by mitochondria because of damage to electron transport chain proteins (Adam-Vizi, 2005; Wang and Michaelis, 2010) 2) compromised antioxidant defense systems leading to an increase in the level, spatial extent, and influence of ROS signaling (Wang et al., 2009) 3) long-term modification of K_ATP_ channel gating and expression (Shen and Johnson, 2012).

If circuit activity is moderately perturbed in the prodromal phase of PD, homeostatic cellular and synaptic plasticity may help to maintain activity and encoding within normal functional limits, and thus delay the expression of motor dysfunction. However, in the 6-OHDA model studied here, NMDAR-dependent STN plasticity appears maladaptive because prevention through NMDAR KD reduced motor dysfunction (Chu et al., 2017). Also consistent with the maladaptive nature of STN cellular plasticity is the demonstration that chemogenetic restoration of autonomous activity ameliorated Parkinsonian motor dysfunction. Therefore, it would be interesting to know when, and to what degree, both cellular and synaptic STN plasticity are engaged during progressive dopamine neuron degeneration. If STN plasticity is maladaptive and contributes to motor dysfunction then it should become progressively extreme as dopamine neurons degenerate. Alternatively, STN plasticity could be maximally engaged prior to the onset of motor symptoms, and circuit dysfunction and symptomatic expression result from increasingly aberrant upstream activity. In this case, chemogenetic restoration of autonomous STN activity may be therapeutic because it opposes pathological patterning of STN activity by the upstream components of the hyperdirect and indirect pathways (Mallet et al., 2008a; Tachibana et al., 2011). Consistent with this concept was the observation that chemogenetic activation of STN neurons reduced the number of synaptically driven spikes both in absolute terms and as a percentage of those generated intrinsically during optogenetic stimulation of cortico-STN inputs at β band frequency. Thus, chemogenetic activation of STN neurons *in vivo* may interrupt the propagation of correlated and coherent activity patterns that have been linked to the expression of Parkinsonian akinesia, bradykinesia, and rigidity (Jenkinson and Brown, 2011; Sanders and Jaeger, 2016; Sharott et al., 2014; Shimamoto et al., 2013). Indeed, STN NMDAR KD, which also ameliorates motor dysfunction, not only prevents loss of autonomous STN activity but also disconnects the indirect pathway through the potent downregulation of the GPe-STN synapses (Chu et al., 2015). Together these studies argue that interrupting the pathological synaptic patterning of STN activity by both the cortex and the GPe may be key to therapy and underlie the beneficial effects of diverse STN manipulations, including lesions (Bergman et al., 1990), optogenetic/pharmacological silencing of STN activity (Levy et al., 2001; Yoon et al., 2014), high-frequency optogenetic stimulation of cortico-STN axon terminals (Gradinaru et al., 2009; Sanders and Jaeger,2016), and direct high-frequency electrical stimulation of the nucleus (Benabid et al., 2009; Wichmann et al., 2018).

## ACKNOWLEDGEMENTS

This study was funded by NIH NINDS grants 2R37 NS041280, P50 NS047085, 5T32 NS041234 and F31 NS090845. Confocal imaging work performed at the Northwestern University Center for Advanced MicROScopy, supported by NCI CCSG grant P30 CA060553. We thank S.Ulrich, D.R.Schowalter and M.Alicea for mouse colony management.

## AUTHOR CONTRIBUTIONS

Conceptualization, E.L.M., M.D.B.; Pilot experiments, E.L.M., K.E.C., J.F.A., M.D.B.; Methodology: E.L.M., H.-Y.C., J.F.A., J.K., D.W., D.J.S., M.D.B.; Formal Analysis/Investigation: E.L.M., H.-Y.C., J.F.A., M.D.B.; Writing – original Draft, E.L.M., M.D.B.; Writing – Review and Editing, E.L.M, H.-Y.C., J.F.A., K.E.C., J.K., D.W., D.J.S., M.D.B.; Visualization, E.L.M, H.-Y.C., J.F.A., D.W., M.D.B.; Supervision, Project Administration, Funding Acquisition: E.L.M, M.D.B.

---

**Figure S1.**
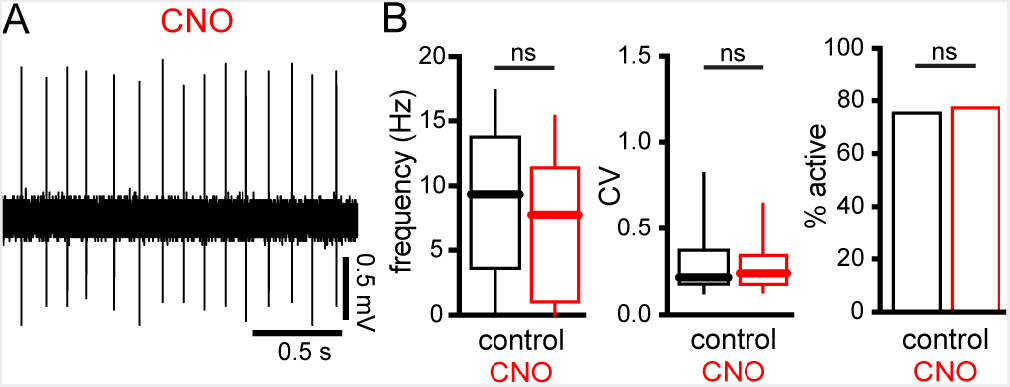
Relates to Figure 2 and Table S2. Chronic SC administration of CNO had no effect on autonomous STN activity in C57BL/6 mice that do not express rM3D(Gs)-mCherry. (A,B) Representative trace (A) and population data (B) demonstrating that administration of CNO according to the schedule in **Figure 2A** had no effect on the frequency, regularity, and incidence of autonomous STN activity firing in C57BL/6 mice. Control refers to the data illustrated in **Figure 1C, D, H.**ns, not significant.

**Figure S2.**
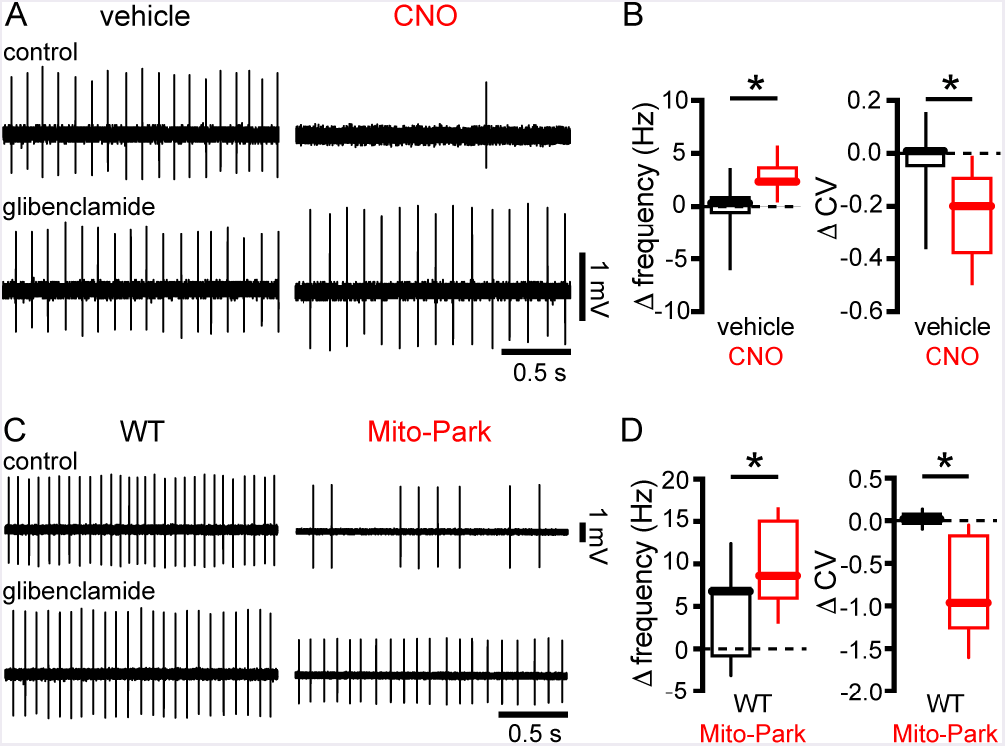
Relates to Figures 4 and S3 and Table S4. Inhibition of **K_ATP_** channels rescued autonomous STN activity in other models of Parkinsonism. (A,B) Inhibition of K_ATP_ channels with glibenclamide rescued the frequency and regularity of autonomous firing in slices from CNO-treated adora2A-rM3D(Gs)-mCherry mice but had minimal effects in slices from vehicle-treated adora2A-rM3D(Gs)-mCherry mice example; B, population data).(C, D) Inhibition of K_ATP_ channels with glibenclamide rescued the frequency and regularity of autonomous firing in slices from 20-week old Mito-Park mice but had minimal effects in age-matched control mice (C, example; D, population data).* p < 0.05.

**Figure S3.**
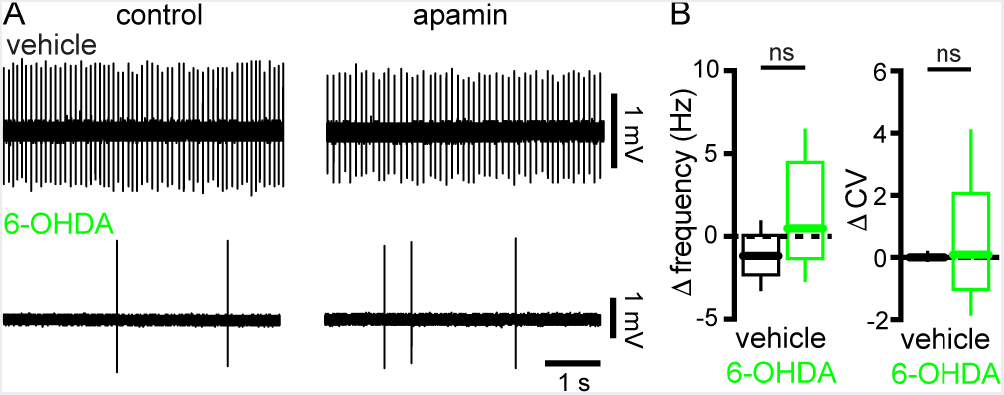
Relates to Figures 4 and S2 and Table S4. Inhibition of SK channels did not rescue autonomous activity in 6-OHDA-injected mice. (A,B) Inhibition of SK channels with 10 nM apamin did not rescue the frequency and regularity of autonomous firing in slices from 6-OHDA-injected mice. Furthermore, the effects of SK channel inhibition were similar in 6-OHDA- and vehicle-injected mice example; B, population data).ns, not significant.

**Table S1.**
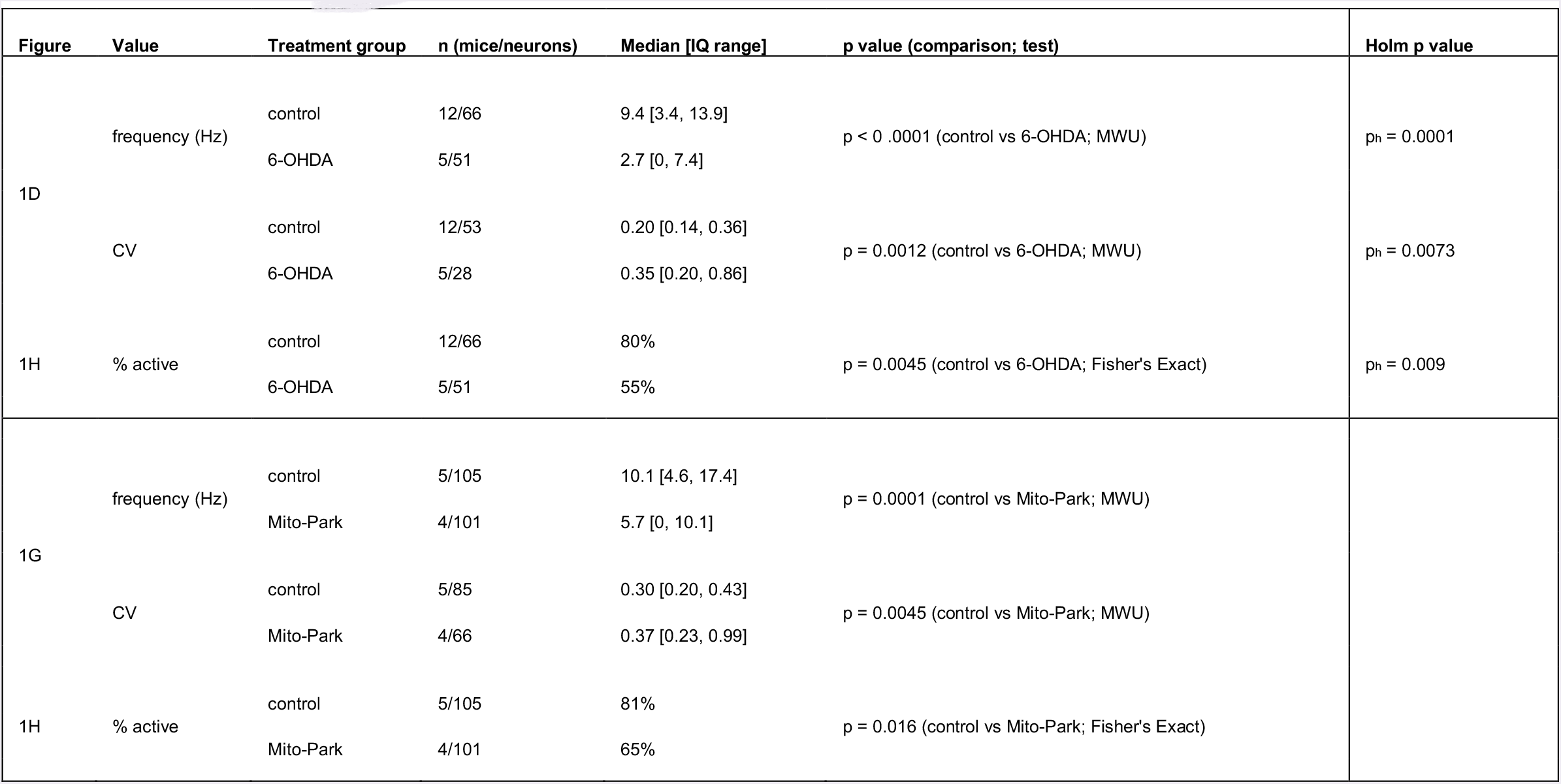
Relates to Figure 1.

| Figure | Value | Treatment group | n (mice/neurons) | Median [IQ range] | p value (comparison; test) | Holm p value |
| --- | --- | --- | --- | --- | --- | --- |
| 1D | frequency (Hz) | control | 12/66 | 9.4 [3.4, 13.9] | p < 0 .0001 (control vs 6-OHDA; MWU) | p<sub>h</sub> = 0.0001 |
|  |  | 6-OHDA | 5/51 | 2.7 [0, 7.4] |  |  |
|  | CV | control | 12/53 | 0.20 [0.14, 0.36] | p = 0.0012 (control vs 6-OHDA; MWU) | p<sub>h</sub> = 0.0073 |
|  |  | 6-OHDA | 5/28 | 0.35 [0.20, 0.86] |  |  |
| 1H | % active | control | 12/66 | 80% | p = 0.0045 (control vs 6-OHDA; Fisher's Exact) | p<sub>h</sub> = 0.009 |
|  |  | 6-OHDA | 5/51 | 55% |  |  |
| 1G | frequency (Hz) | control | 5/105 | 10.1 [4.6, 17.4] | p = 0.0001 (control vs Mito-Park; MWU) |  |
|  |  | Mito-Park | 4/101 | 5.7 [0, 10.1] |  |  |
|  | CV | control | 5/85 | 0.30 [0.20, 0.43] | p = 0.0045 (control vs Mito-Park; MWU) |  |
|  |  | Mito-Park | 4/66 | 0.37 [0.23, 0.99] |  |  |
| 1H | % active | control | 5/105 | 81% | p = 0.016 (control vs Mito-Park; Fisher's Exact) |  |
|  |  | Mito-Park | 4/101 | 65% |  |  |

**Table S2.** Relates to Figures 2 and S1.

| Figure | Value | Treatment group | n (animals/neurons) | Median [IQ range] | p value (comparison; test) | Holm p value |
| --- | --- | --- | --- | --- | --- | --- |
| 2D | frequency (Hz) | A2A/vehicle | 3/67 | 10.1 [3.9, 16.6] | p < 0.0001 (veh vs CNO; MWU) |  |
|  |  | A2A/CNO | 3/67 | 0 [0, 3.0] |  |  |
|  | CV | A2A/vehicle | 3/63 | 0.16 [0.10, 0.39] | p = 0.0038 (veh vs CNO; MWU) |  |
|  |  | A2A//CNO | 3/29 | 0.45 [0.16, 1.1] |  |  |
|  | % active | A2A//vehicle | 3/67 | 94% | p < 0.0001 (veh vs CNO; Fisher's Exact) |  |
|  |  | A2A//CNO | 3/67 | 43% |  |  |
| 2E, F | frequency (Hz) | A2A MFB veh/SC veh | 3/60 | 7.3 [0.33, 13.7] | p < 0.0001 (veh/veh vs MFB veh/SC CNO; MWU) | ph = 0.0001 |
|  |  | A2A MFB veh/SC CNO | 3/58 | 0.61 [0, 5.1] | p = 0.0024 (veh/veh vs 6-OHDA/veh; MWU) | ph = 0.0048 |
|  |  | A2A MFB 6-OHDA/SC veh | 3/61 | 3.5 [0, 7.8] | p = 0.50 (6-OHDA/veh vs 6-OHDA/CNO; MWU) | ph = 0.50 |
|  |  | A2A MFB 6-OHDA/SC CNO | 3/60 | 0.60 [0, 7.4] |  |  |
|  | CV | A2A MFB veh/SC veh | 3/47 | 0.27 [0.15, 0.50] | p = 0.0003 (veh/veh vs veh/ CNO; MWU) | ph = 0.0010 |
|  |  | A2A MFB veh/SC CNO | 3/33 | 0.70 [0.34, 1.3] | p = 0.42 (veh/veh vs 6-OHDA/veh; MWU) | ph = 0.85 |
|  |  | A2A MFB 6-OHDA/SC veh | 3/37 | 0.35 [0.17, 1.0] | p = 0.95 (6-OHDA/veh vs 6-OHDA/CNO; MWU) | ph = 0.95 |
|  |  | A2A MFB 6-OHDA/SC CNO | 3/33 | 0.38 [0.15, 0.94] |  |  |
|  | % active | A2A MFB veh/SC veh | 3/60 | 80% | p = 0.0012 (veh/veh vs veh/ CNO; MWU) | ph = 0.0036 |
|  |  | A2A MFB veh/SC CNO | 3/58 | 58% | p = 0.0050 (veh/veh vs 6-OHDA/veh; MWU) | ph = 0.01 |
|  |  | A2A MFB 6-OHDA/SC veh | 3/61 | 61% | p = 0.47 (6-OHDA/veh vs 6-OHDA/CNO; MWU) | ph = 0.47 |
|  |  | A2A MFB 6-OHDA/SC CNO | 3/60 | 55% |  |  |
| S1B | frequency (Hz) | C57/CNO | 2/37 | 7.8 [0.9, 11.6] | p = 0.22 (Fig 1D control vs C57/CNO; MWU) | ph = 0.66 |
|  | CV | C57/CNO | 2/28 | 0.24 [0.16, 0.35] | p = 0.38 (Fig 1D control vs C57/CNO; MWU) | ph = 0.75 |
|  | % active | C57/CNO | 2/37 | 78% | p > 0.9999 (Fig 1H control vs C57/CNO; Fisher's Exact) | ph = 1 |

**Table S3.**
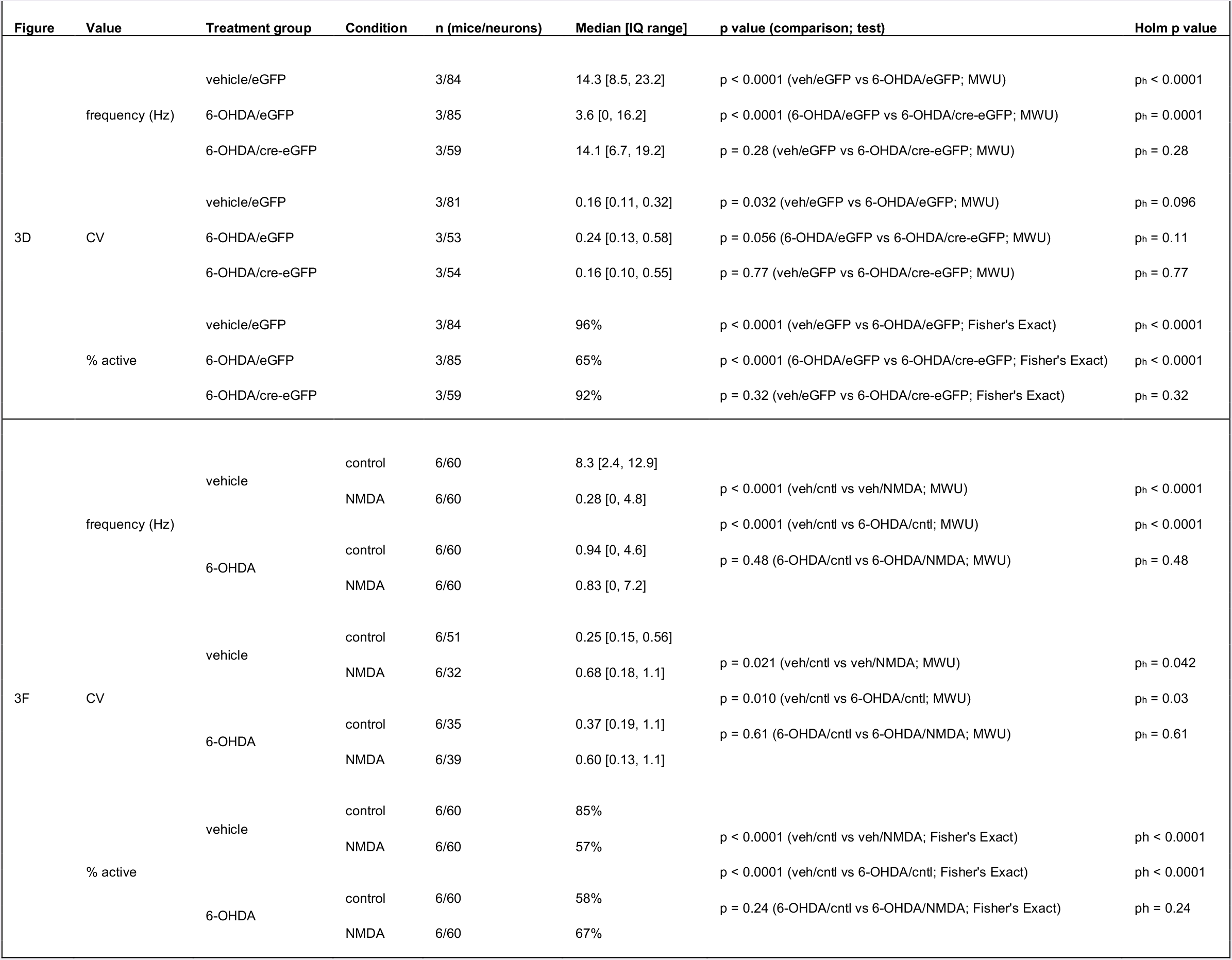
Relates to Figure 3.

**Table S4.**
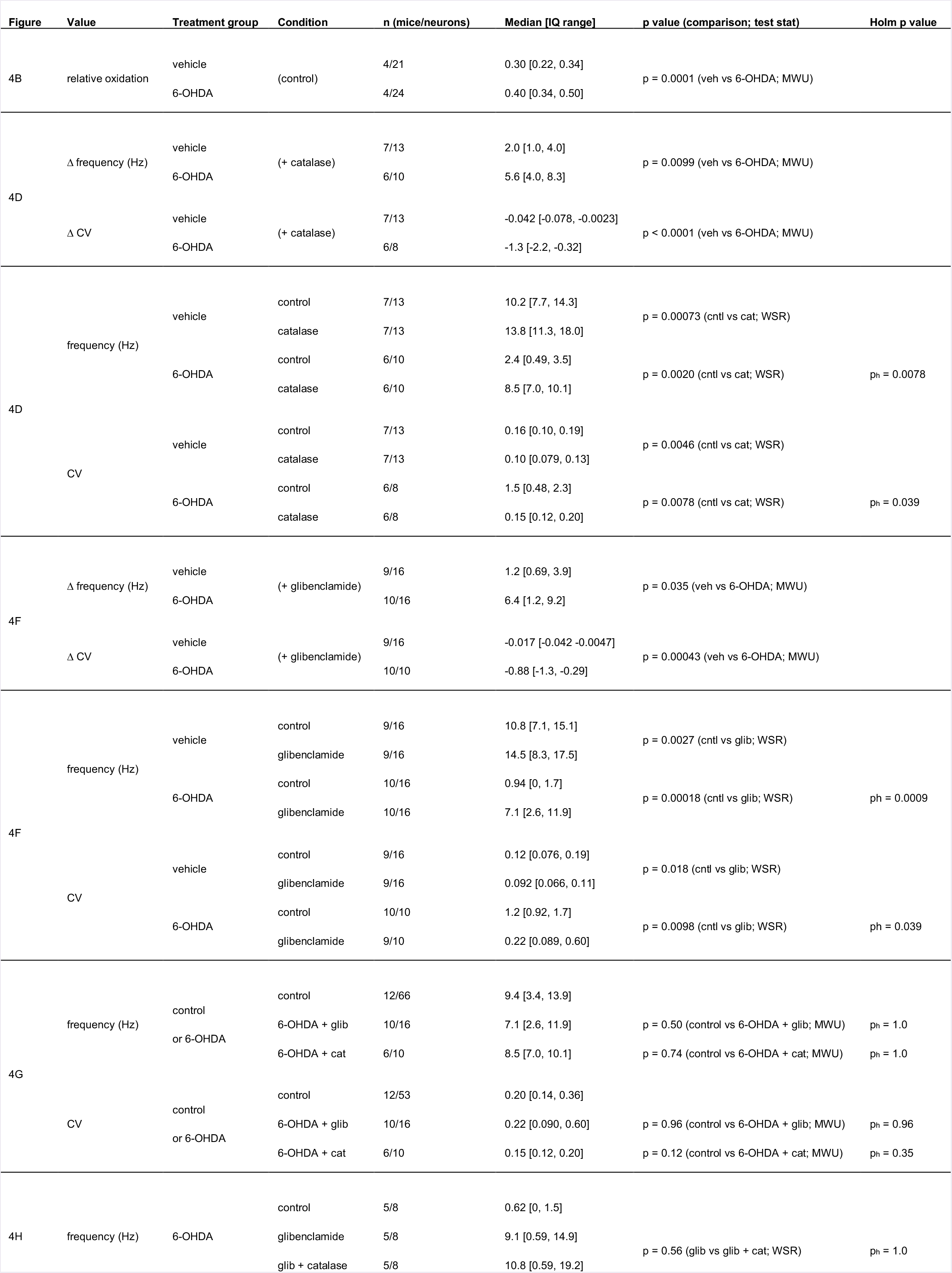
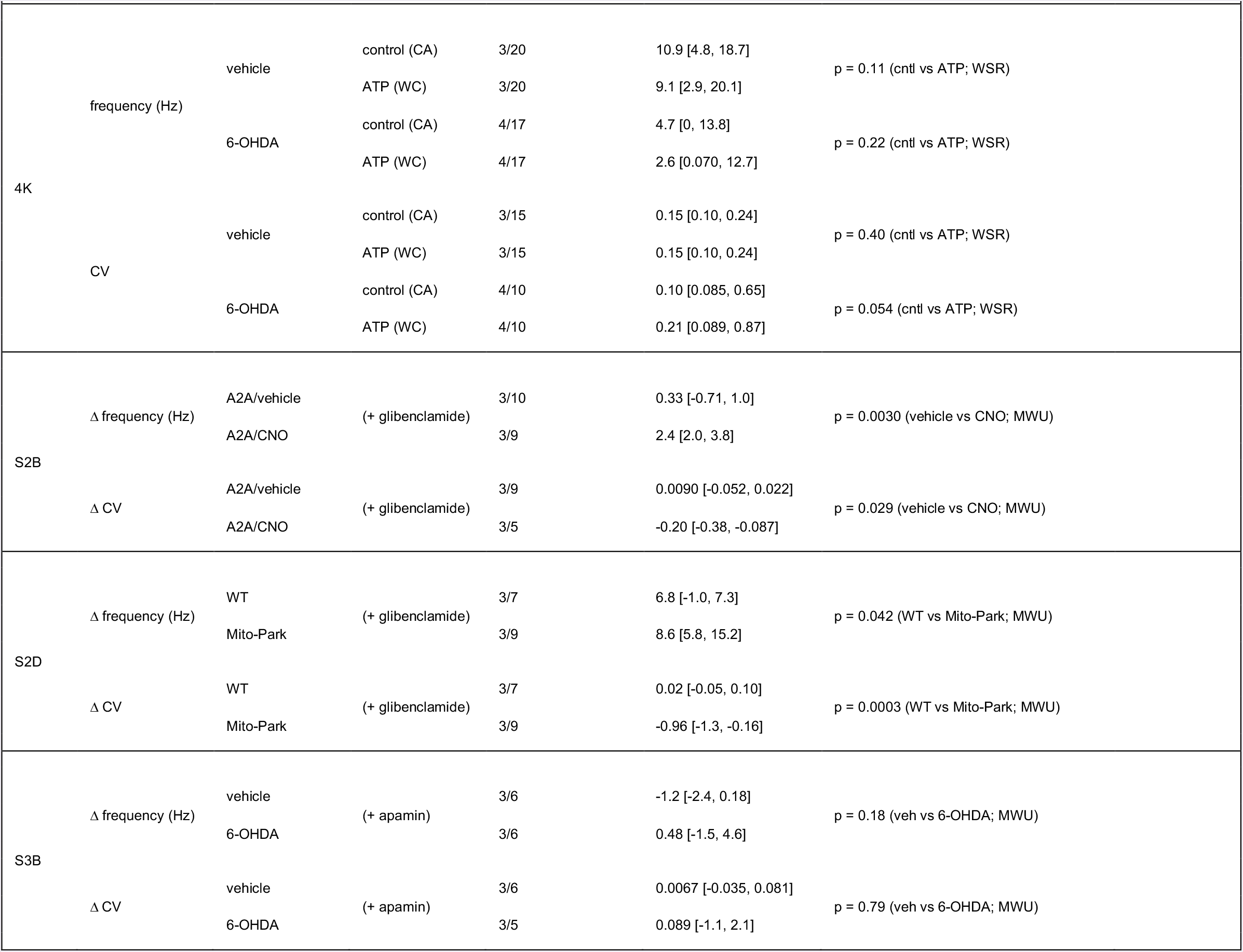
Relates to Figures 4, S2, and S3.

**Table S5.**
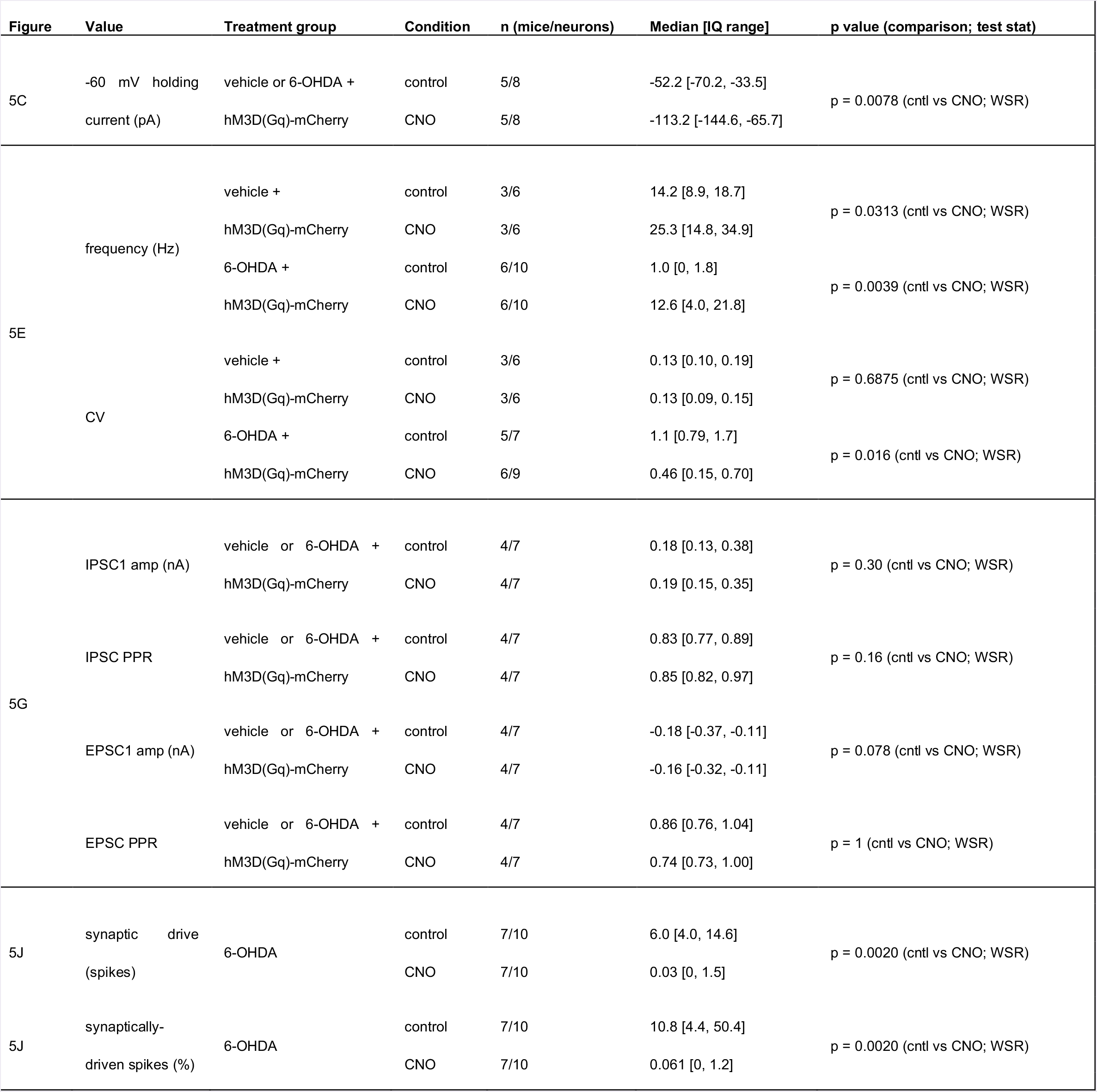
Relates to Figure 5.

| Figure | Value | Treatment group | Condition | n (mice/neurons) | Median [IQ range] | p value (comparison; test stat) |
| --- | --- | --- | --- | --- | --- | --- |
| 5C | -60 mV holding | vehicle or 6-OHDA + | control | 5/8 | -52.2 [-70.2, -33.5] | p = 0.0078 (cntl vs CNO; WSR) |
|  | current (pA) | hM3D(Gq)-mCherry | CNO | 5/8 | -113.2 [-144.6, -65.7] |  |
| 5E | frequency (Hz) | vehicle + | control | 3/6 | 14.2 [8.9, 18.7] | p = 0.0313 (cntl vs CNO; WSR) |
|  |  | hM3D(Gq)-mCherry | CNO | 3/6 | 25.3 [14.8, 34.9] |  |
|  |  | 6-OHDA + | control | 6/10 | 1.0 [0, 1.8] | p = 0.0039 (cntl vs CNO; WSR) |
|  |  | hM3D(Gq)-mCherry | CNO | 6/10 | 12.6 [4.0, 21.8] |  |
|  | CV | vehicle + | control | 3/6 | 0.13 [0.10, 0.19] | p = 0.6875 (cntl vs CNO; WSR) |
|  |  | hM3D(Gq)-mCherry | CNO | 3/6 | 0.13 [0.09, 0.15] |  |
|  |  | 6-OHDA + | control | 5/7 | 1.1 [0.79, 1.7] | p = 0.016 (cntl vs CNO; WSR) |
|  |  | hM3D(Gq)-mCherry | CNO | 6/9 | 0.46 [0.15, 0.70] |  |
| 5G | IPSC1 amp (nA) | vehicle or 6-OHDA + | control | 4/7 | 0.18 [0.13, 0.38] | p = 0.30 (cntl vs CNO; WSR) |
|  |  | hM3D(Gq)-mCherry | CNO | 4/7 | 0.19 [0.15, 0.35] |  |
|  | IPSC PPR | vehicle or 6-OHDA + | control | 4/7 | 0.83 [0.77, 0.89] | p = 0.16 (cntl vs CNO; WSR) |
|  |  | hM3D(Gq)-mCherry | CNO | 4/7 | 0.85 [0.82, 0.97] |  |
|  | EPSC1 amp (nA) | vehicle or 6-OHDA + | control | 4/7 | -0.18 [-0.37, -0.11] | p = 0.078 (cntl vs CNO; WSR) |
|  |  | hM3D(Gq)-mCherry | CNO | 4/7 | -0.16 [-0.32, -0.11] |  |
|  | EPSC PPR | vehicle or 6-OHDA + | control | 4/7 | 0.86 [0.76, 1.04] | p = 1 (cntl vs CNO; WSR) |
|  |  | hM3D(Gq)-mCherry | CNO | 4/7 | 0.74 [0.73, 1.00] |  |
| 5J | synaptic drive<br>(spikes) | 6-OHDA | control | 7/10 | 6.0 [4.0, 14.6] | p = 0.0020 (cntl vs CNO; WSR) |
|  |  |  | CNO | 7/10 | 0.03 [0, 1.5] |  |
| 5J | synaptically-driven spikes (%) | 6-OHDA | control | 7/10 | 10.8 [4.4, 50.4] | p = 0.0020 (cntl vs CNO; WSR) |
|  |  |  | CNO | 7/10 | 0.061 [0, 1.2] |  |

**Table S6.** Relates to Figure 6.

| Figure | Value | Treatment group | Condition | n (mice/sites) | Median [IQ range] | p value (comparison; test stat) |
| --- | --- | --- | --- | --- | --- | --- |
| 6C | density of cfos+ neurons in STN (10<sup>3</sup>.mm<sup>-3</sup>) | vehicle + | vehicle | 3/18 | 0 [0, 8.9] | p < 0.0001 (vehicle vs CNO; MWU) |
|  |  | hM3D(Gq) | CNO | 3/18 | 100.4 [43.5, 144.1] |  |
|  |  | 6-OHDA + | vehicle | 3/18 | 0 [0, 0] | p < 0.0001 (vehicle vs CNO; MWU) |
|  |  | hM3D(Gq) | CNO | 3/18 | 129.5 [79.81, 180.6] |  |
| 6F | density of cfos+ PV+ neurons in SNr (10<sup>3</sup>.mm<sup>-3</sup>) | vehicle + | vehicle | 3/18 | 0 [0, 0] | p < 0.0001 (vehicle vs CNO; MWU) |
|  |  | hM3D(Gq) | CNO | 3/18 | 37.1 [0, 44.8] |  |
|  |  | 6-OHDA + | vehicle | 3/18 | 0 [0, 33.4] | p < 0.0001 (vehicle vs CNO; MWU) |
|  |  | hM3D(Gq) | CNO | 3/18 | 86.3 [44.1, 105.4] |  |

**Table S7.**
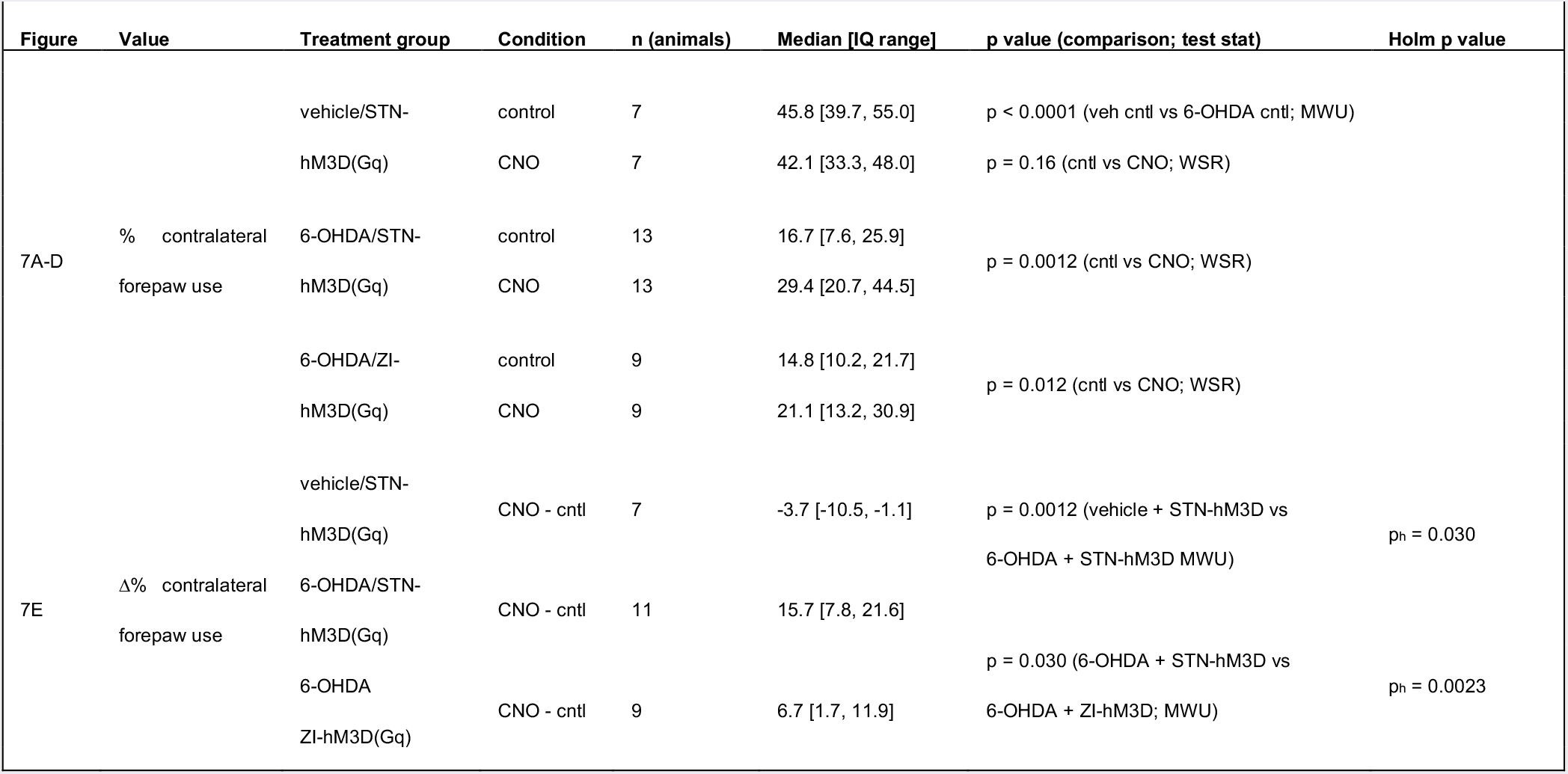
Relates to Figure 7.

Relates to Experimental Procedures.

**Table S8.** Relates to Experimental Procedures.
| Figure | Value | Treatment group | n (animals) | Median [IQ range] | p value (comparison; test stat) |
| --- | --- | --- | --- | --- | --- |
| text | age at experiment (days) | All groups | 190 | 81 [70, 97] |  |
|  |  | C57BL/6 | 152 | 76 [70, 92] |  |
|  |  | adora2a-rM3D(s) | 18 | 125 [87, 205] |  |
|  |  | mitoPark | 9 | 138 [130, 170] |  |
|  |  | GRIN1 | 9 | 89 [82, 198] |  |
|  |  | Gabrr3-cre | 2 | 68 |  |
| text | time after surgery (days) | All groups | 173 | 20 [16, 22] |  |
|  |  | C57BL/6 | 150 | 20 [16, 22] |  |
|  |  | adora2a-rM3D(s) | 12 | 22 [21, 30] |  |
|  |  | GRIN1 | 9 | 16 [16, 22] |  |
|  |  | Gabrr3-cre | 2 | 15 |  |
| T | % striatal TH remaining in | vehicle | 62 | 96.5 [86.9, 113.5] | p < 0.0001 |
| ext | ipsilateral hemisphere | 6-OHDA | 98 | 0.5 [0, 7.0] |  |

## STAR METHODS

### CONTACT FOR REAGENT AND RESOURCE SHARING

Further information and requests for resources and reagents should be directed to the Lead Contact, Mark Bevan <u></u>.

### EXPERIMENTAL MODEL AND SUBJECT DETAILS

Experiments were performed using naïve adult male C57BL/6 (RRID:IMSR_JAX:000664; age = 76, 70-92 days; n = 152), B6. Cg-*Tfam^tm1.1Ncdl^/J* (RRID:IMSR_JAX:026123) X B6. SJL-*Slc6a3;^tm1.1(cre)Bkmn^/J* (RRID:IMSR_JAX:006660; age = 138, 130-170 days, n = 9), B6.129S4-*Grin1^tm2Stl^/J* (RRID:IMSR_JAX:005246; age = 89, 82-198 days; n = 9), B6. Cg-Tg(Adora2a-Chrm3*,-mCherry)AD6Blr/J (RRID:IMSR_JAX:017863; age = 125, 87-205 days; n = 18), and Tg(Gabrr3-cre)KC112Gsat (RRID:MGI:5528400; age = 68 days; n = 2) mice (**Table S8**) according to IACUC and NIH guidelines for the care and use of animals. Mice were housed singly or with up to 4 other littermates per cage under a 14-hour light, 10-hour dark cycle with food and water *ad libitum*, and were inspected daily by animal care technicians, veterinarians, and laboratory staff. Where possible littermates were assigned evenly to different treatment groups.

## METHOD DETAILS

### Surgery

6-OHDA, vehicle, and viral vectors were injected stereotaxically (NeuROStar, Tubingen, Germany) under (1-2%) isoflurane anesthesia (Smiths Medical ASD, Inc., Dublin, oH, USA).6-OHDA (3-4 mg/ml) or vehicle was injected unilaterally into the medial forebrain bundle (MFB; from Bregma: AP -0.7 mm, ML +1.2 mm, DV +4.7 mm; 1.5 μl) to lesion midbrain dopamine neurons or control for injection, respectively. Pargyline (50 mg/kg) and desipramine (25 mg/kg) were used to increase the potency and specificity of 6-OHDA, respectively. In C57BL/6 mice AAV expressing hSyn-ChR2(H134R)-eYFP (AAV9.hSyn.hChR2(H134R)-eYFP. WPRE.hGH; 1 × 10^13^ GC/ml; University of Pennsylvania Viral Vector Core) was injected at 3 sites in primary motor cortex (AP +0.6 mm, +1.2 mm, and +1.8 mm, ML +1.5 mm, DV +1.0 mm; 300 nl per site), or AAV expressing CMV-MTS-roGFP (AAV9-CMV-mito-roGFP-SV40; 2.5 × 10^12^ GC/ml; Virovek Inc., Hayward, CA), or hSyn-HA-hM3D(Gq)-IRES-mCitrine (AAV2/5-hSyn-HA-hM3D(Gq)-IRES-mCitrine; 2 × 10^12^ GC/ml) or hSyn-hM3D(Gq)-mCherry (AAV8-hSyn-hM3D(Gq)-mCherry; 0.5-3 × 10^12^ GC/ml; UNC Vector Core, Chapel Hill, NC or Addgene, Cambridge, MA) was injected in the STN (AP -2.06 mm, ML +1.4 mm, DV +4.45 mm; 300-500 nl), ipsilateral to the MFB injection. In *Grin1^lox/lox^* mice AAV expressing eGFP (AAV9-hSyn-eGFP-WPRE-bGH; 2 × 10^12^ GC/ml; University of Pennsylvania Viral Vector Core) or cre-eGFP (AAV9-hSyn-HI-eGFP-Cre-WPRE-SV40; 2 × 10^12^ GC/ml; University of Pennsylvania Viral Vector Core) was injected in the STN (AP -2.06 mm, ML +1.4 mm, DV +4.45 mm; 150-300 nl) ipsilateral to the MFB injection. Gabrr3-cre mice, in which there is cre-recombinase expression in STN neurons and minimal expression in adjacent structures, were injected with AAV expressing cre-dependent hSyn-DIo-hM3D(Gq)-mCherry (AAV8-hSyn-DIo-hM3D(Gq)-mCherry; 2-3 × 10^12^ GC/ml; UNC Vector Core, Chapel Hill, NC) in the STN. Finally, a subset of C57BL/6 mice were injected with AAV8-hSyn-hM3D(Gq)-mCherry dorsal to the STN (AP -2.06, ML +1.4, DV +3.75) to test the behavioral effects of hM3D(Gq) activation of structures adjacent to the STN. Electrophysiological, imaging and behavioral measurements were made 20, 16-22 days (n = 173) after surgery.

### Electrophysiology

Animals were first anesthetized through IP injection of ketamine/xylazine (87/13 mg/kg) and then transcardially perfused with ice-cold sucROSe-based artificial cerebro-spinal fluid (sACSF: 230 mM sucROSe, 2.5 mM KCl, 1.25 mM NaH_2_Po4, 0.5 mM CaCl_2_, 10 mM MgSo_4_, 10 mM glucose, and 26 mM NaHCo_3_), equilibrated with 95% o2 and 5% Co_2_. The brain was then removed and sectioned at 250 μm in the parasagittal plane at 30 μm/s in a chamber containing ice-cold sACSF using a vibratome (VT1200S; Leica MicROSystems Inc., Buffalo Grove, IL, USA). Cut slices were transferred to ACSF (126 mM NaCl, 2.5 mM KCl, 1.25 mM NaH_2_Po_4_, 2 mM CaCl_2_, 2 mM MgSo_4_, 10 mM glucose, 26 mM NaHCo_3_, 1 mM sodium pyruvate, and 5 μM L-glutathione), equilibrated with 95% o_2_ and 5% Co2 at 35¼C for 30 minutes and then held in ACSF at room temperature until recording.

In the recording chamber slices were perfused at a rate of 4-5 ml/min with synthetic interstitial fluid (SIF: 126 mM NaCl, 3 mM KCl, 1.25 mM NaH_2_PO_4_, 1.6 mM CaCl_2_, 1.5 mM MgSO_4_, 10 mM glucose and 26 mM NaHCO_3_), equilibrated with 95% o2 and 5% Co2 at 35¼C. A micROScope (Axioskop FS2 micROScope (Carl Zeiss, oberkochen, Germany) equipped with a LUMPlanFl/IR 60 X 0.9 NA objective (olympus, Tokyo, Japan) or a BX51WI micROScope (olympus) equipped with a UIS2 LUMPFL 60 × 0.9 NA objective (olympus)) employing infrared Dodt Gradient Contrast illumination (Luigs & Neumann, Ratingen, Germany) was used to visualize cell bodies for patch clamp recording.590 nm LED illumination (Cairn Instruments, Faversham, Kent, UK) was used to identify neurons expressing hM3D(Gq)-mCherry and 890 nm 2-photon illumination (Mira 900F and G8 532 nm oPSL pump laser, Coherent Inc., Santa Clara, USA; as for MTS-roGFP imaging, detailed below) was used to identify neurons expressing eGFP or cre-eGFP.

Recordings were obtained using computer-controlled manipulators (Luigs & Neumann) and a Multiclamp 700B amplifier, and a Digidata 1440A digitizer controlled by PClamp10 (Molecular Devices, Sunnyvale, CA, USA). Signals were low-pass filtered online at 10 kHz and sampled at 50 kHz. Cell and electrode capacitances, series resistance, and junction potentials were compensated electronically. Recordings utilized boROSilicate glass pipettes (Warner Instruments, Hamden, CT, USA) pulled using a P-97 Flaming/Brown Micropipette Puller (Sutter Instruments, Novato, CA, USA). Loose-seal cell-attached recordings were made with 3-5 MΩ impedance pipettes containing HEPES-buffered SIF solution (HBS: 140 mM NaCl, 23 mM glucose, 15 mM HEPES, 3 mM KCl, 1.5 mM MgCl_2_, 1.6 mM CaCl_2_; pH 7.2 with NaoH; 300–310 mosm/L). Cell-attached recordings were excluded if the membrane became disrupted. Whole-cell voltage clamp recordings were made using 4-5 MΩ impedance pipettes filled with a K-gluconate-based internal solution (140 mM K-gluconate, 3.8 mM NaCl, 1 mM MgCl2, 10 mM HEPES, 0.1 mM Na_4_-EGTA, 0.4 mM Na_3_GTP, and 2 mM Mg_1.5_ATP). Voltage clamp recordings were excluded if series resistance changed by more than 20% during the course of recording. Whole-cell current clamp recordings were made using 12-15 MΩ impedance pipettes filled with KCH3SO4-based internal solution (130 mM KCH_3_So_4_, 3.8 mM NaCl, 1 mM MgCl2, 10 mM HEPES, 5 mM phosphocreatine, 0.1 mM Na4-EGTA, 0.4 mM Na_3_GTP, and 2 mM Mg_1.5_ATP).

Optogenetic stimulation was delivered via the objective lens using a 470 nm light emitting diode (OptoLED; Cairn Research, Faversham, Kent, UK). Cortico-STN transmission was evoked by 1 ms duration optogenetic stimulation of cortico-STN axon-terminals. EPSCs and IPSCs were also evoked by electrical stimulation of the internal capsule bordering the ROStro-ventral STN using a constant current isolator (A365R, World Precision Instruments, Sarasota, FL). The poles of stimulation were selected from a custom-built matrix of 10 stimulation electrodes (MX52CBWMB2, Frederick Haer, Bowdoin, ME).

Autonomous activity was typically recorded in the presence of 20 μM DNQX, 50 μM D-APV, 10 μM SR-95531, and 2 μM CGP55845 (Abcam, Cambridge, MA, USA) to antagonize AMPARs, NMDARs, GABAaRs and GABAbRs, respectively. DNQX and D-APV were excluded when the response of STN neurons to optogenetic cortico-STN stimulation or electrically evoked EPSCs were studied. SR-95531 and CGP55845 were excluded when electrically evoked IPSCs were studied. The effect of catalase (polyethylene glycol-catalase; 250 U/mL; Sigma-Aldrich), glibenclamide (100 nM; Sigma-Aldrich), apamin (10 nM; Sigma-Aldrich), or CNO (10-100 μM; Sigma-Aldrich) on firing was also studied.

### 2-photon imaging of mito-roGFP

Brain slices were prepared from mice expressing the mitochondrially-targeted redox-sensitive probe MTS-roGFP in STN neurons, as for electrophysiology. MTS-roGFP-expressing neurons were imaged at 890 nm with 76 MHz pulse repetition and ~250 fs pulse duration at the sample plane. Two-photon excitation was provided by a G8 oPSL pumped Mira 900 F laser (Coherent, Santa Clara, CA, USA). Sample power was regulated by a Pockels cell electro-optic modulator (model M350-50-02-BK, Con optics, Danbury, CT, USA). Images were acquired using an Ultima 2 P system running *PrairieView* 5.3 (Bruker Nano Fluorescence MicROScopy, Middleton, WI, USA) and a BX51WI micROScope (olympus, Tokyo, Japan) with a 60 × 0.9 NA objective (UIS2 LUMPFL; olympus). MTS-roGFP fluorescence and simultaneous laser-scanned Dodt contrast images were acquired from each field of STN neurons in the presence of 20 μM DNQX, 50 μM APV, 10 μM SR-95531, and 2 μM CGP55845 under baseline conditions, following the addition of 2 mM dithiothrietol to fully reduce the tissue, and following the addition of 200 μM aldrithiol to fully oxidize the tissue (Atherton et al., 2016). Average fluorescence intensities of mitochondria in STN neurons under baseline, reduced, and oxidized conditions were quantified using Image J (NIH, Bethesda, MD, USA). Baseline mitochondrial oxidation was then expressed relative to that under conditions of minimum and maximum oxidation.

### Behavioral testing

To assess forelimb usage during vertical exploration, animals were placed in a 600 ml cylindrical glass beaker (9.5 cm diameter; 12 cm height) and imaged using an HD digital camcorder recording at 60 fps (VIXIA HF R40 Full HD Camcorder; Canon, Melville, NY, USA). Forelimb placements on the vertical walls of the beaker were counted manually during video review (Schallert et al., 2000). Placement of the left and right forepaw were considered simultaneous (“both”) if they occurred within 3 frames of each other (i.e.< 33 ms apart). Mice expressing hM3D(Gq)-mCherry were tested ~ 20-60 minutes following SC injection of vehicle or 1 mg/kg clozapine-n-oxide (CNO) or in the absence of an injection.

### Histology

In order to study the expression of ChR2(H134)-eYFP, cre-eGFP, eGFP, rM3D(Gs)-mCherry, MTS-roGFP, hM3D(Gq)-mCherry, c-fos, NeuN, PV, and TH brain tissue was first fixed in 4% paraformaldehyde in 0.1 M phosphate buffer, pH7.4. Tissue prepared for electrophysiology or MTS-roGFP imaging was immersion-fixed. Mice used for behavioral analyses were perfused-fixed under deep anesthesia (IP: 87 mg/kg ketamine, 13 mg/kg xylazine). Tissue was held in fixative at 4°C for at least 12 hours before rinsing in phosphate buffered saline (PBS; 0.05 M; pH 7.4). Tissue exceeding 250 μm in thickness was re-sectioned at 70 μm using a vibratome (Leica VT1000S; Leica MicROSystems Inc.). Immunohistochemical detection of c-fos, NeuN, PV, and TH was carried out in PBS containing 0.5% Triton X-100 (Sigma-Aldrich, St. Louis, Mo, USA) and 2% normal donkey serum (Jackson ImmunoResearch, West Grove, PA, USA). Tissue was incubated in primary antibodies for 48-72 hours at 4¼C or overnight at RT (rabbit anti-c-fos: 1:500 dilution; cat# 2250S; Cell Signaling Technology, Danvers, MA; mouse anti-NeuN: 1:200 dilution; cat# MAB377; EMD Millipore, Darmstadt, Germany; guinea pig anti-PV: 1:1,000; cat# 195 004; Synaptic Systems, Gottingen, Germany; mouse anti-TH: 1:500 dilution; cat# MAB318; EMD Millipore), washed in PBS, and then incubated in their respective secondary antibodies (donkey anti-rabbit Alexa Fluor 488; dilution 1:250; cat # 711-545-152; Jackson ImmunoResearch; donkey anti-mouse Alexa Fluor 488; dilution 1:250; cat # 715-545-150; Jackson ImmunoResearch; donkey anti-mouse Alexa Fluor 594; dilution 1:250; cat # 715-585-150; Jackson ImmunoResearch; donkey anti-guinea pig Alexa Fluor 647; dilution 1:250; cat # 706-605-148; Jackson ImmunoResearch) for 90-120 minutes at room temperature before a final wash with PBS. All tissue was mounted on glass slides using ProLong Diamond Antifade Reagent (ThermoFisher Scientific, Waltham, MA, USA) and coverslipped. Sections were imaged using an Axioskop 2 micROScope (Carl Zeiss) equipped with a Neurolucida system (MBF Bioscience, Williston, VT, USA) and/or a confocal laser scanning micROScope (A1R; Nikon, Melville, USA).

The densities of c-fos-positive neurons were assessed using NIH ImageJ and quantified using the optical dissector method (West, 1999). Confocal images were collected using a 60X oil immersion Plan Apo objective lens on a Nikon A1R confocal micROScope (NA=1.4, 0.205 micron/pixel; Nikon, Melville, USA). Sample sites were chosen using a grid (frame size, 50 X 50 μm) superimposed randomly on each image stack. Stereological counting commenced and was terminated at an optical section 5 μm and 21 μm below the slice surface, respectively. Look-up and reference planes were separated by 2 μm. Dopaminergic innervation was assessed from tyROSine hydroxylase immunoreactivity, as described previously (Fan et al., 2012). In the 62 vehicle-injected and 98 6-OHDA-injected mice used in this study ipsilateral striatal TH immunoreactivities were 96.5, 86.9113.5% and 0.5, 0-7% of the contralateral hemisphere, respectively (**Table S8**; p < 0.05).

## QUANTIFICATION AND STATISTICAL ANALYSIS

Only data that were compromised by technical failure (e.g.failure of viral expression or immunohistochemical detection) were excluded. Each dataset, together with the associated number and type of observations, and the statistical tests that were applied are summarized in **Supplemental Tables 1-7**. In addition, paired data points or box-and-whisker plots displaying medians and inter-quartile and 10-90% ranges calculated in Prism 7 (GraphPad Software, Inc., La Jolla, CA) are displayed in their respective figures to reflect the distribution and central tendency of the data. To minimize assumptions concerning the distribution of the data, exact, non-parametric (two-sided) statistics were used throughout: the Mann-Whitney U (MWU) test for unpaired data, the Wilcoxon signed rank (WSR) test for paired comparisons, and Fisher’s exact test for contingency analyses. To ensure our experiments were adequately powered sample sizes for MWU and WSR tests were estimated to achieve a minimum of 80% power. For the MWU test, a 50^th^ percentile difference in median requires 10 samples. For the WSR test, if all pairs exhibit the same direction of change 6-10 observations are required. Exact p values were calculated using the fisher. test and the Wilcox.exact function (exactRankTests package) in R (http://www.r-project.org). Where datasets were subjected to multiple comparisons the p-value was adjusted to maintain the family-wise error rate at 0.05 using the Holm-Bonferroni method (notated ph; (Holm, 1979)).

## DATA AND SOFTWARE AVAILABILITY

Data are available from the authors upon request and will be deposited at https://osf.io/dashboard

---

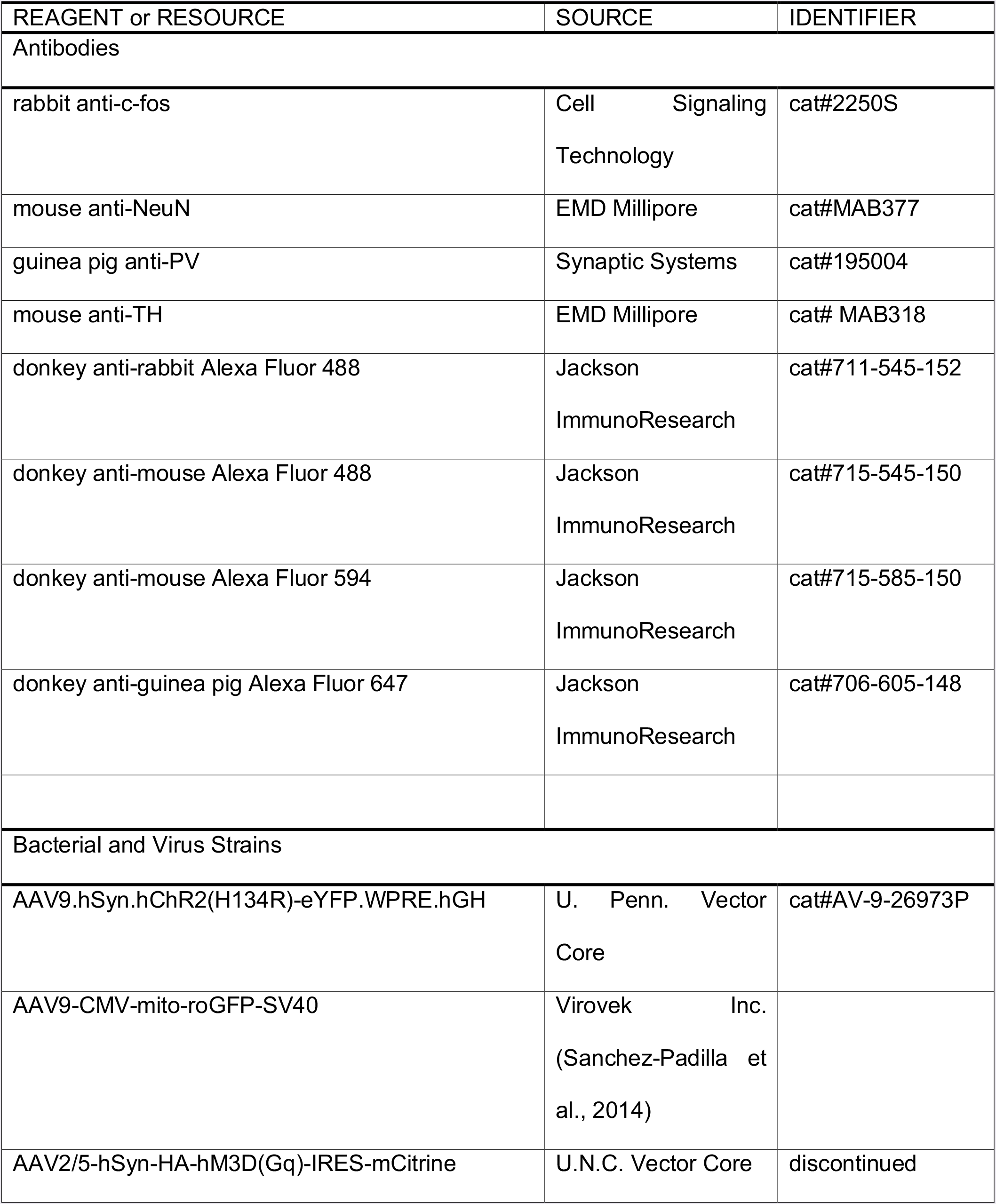
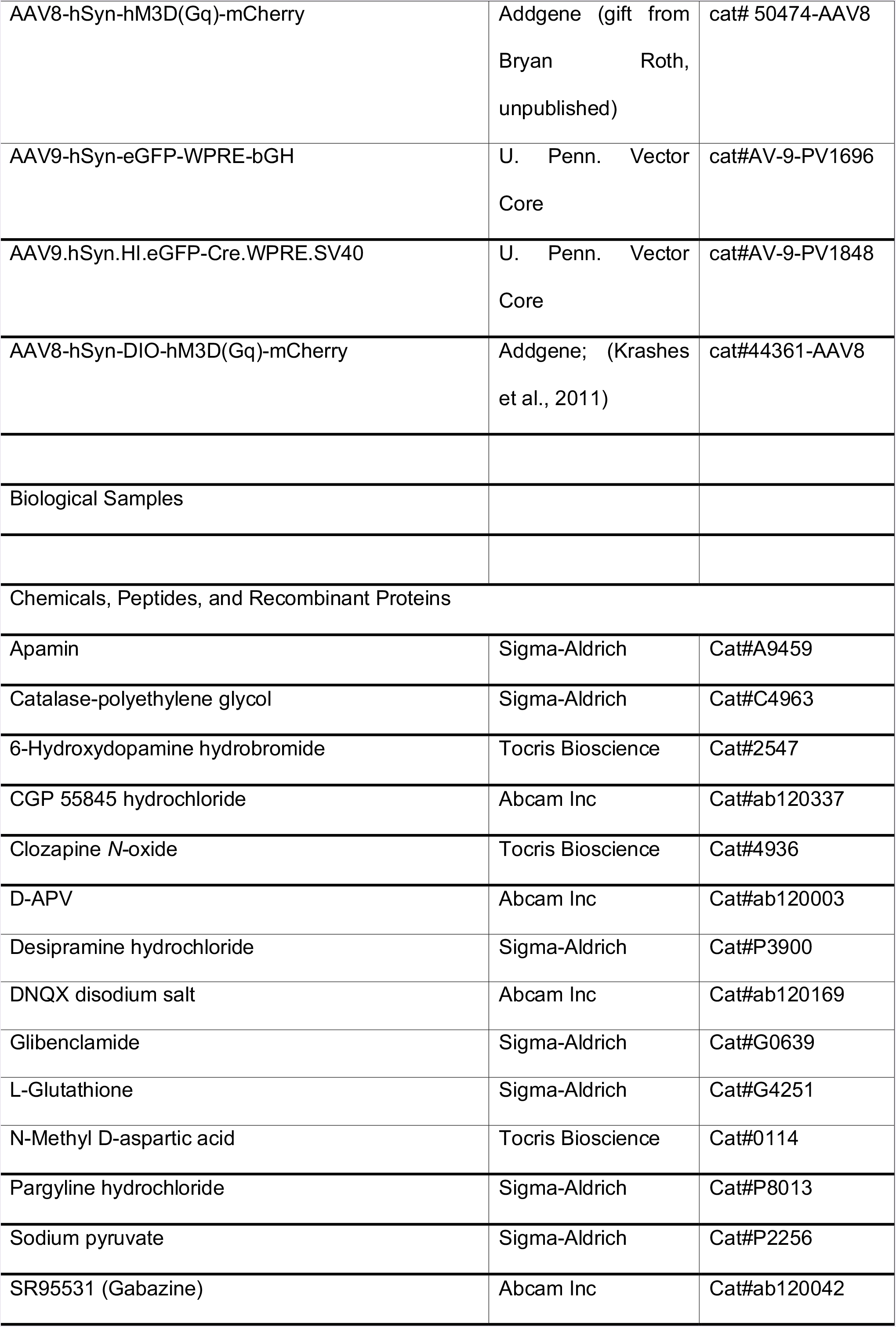
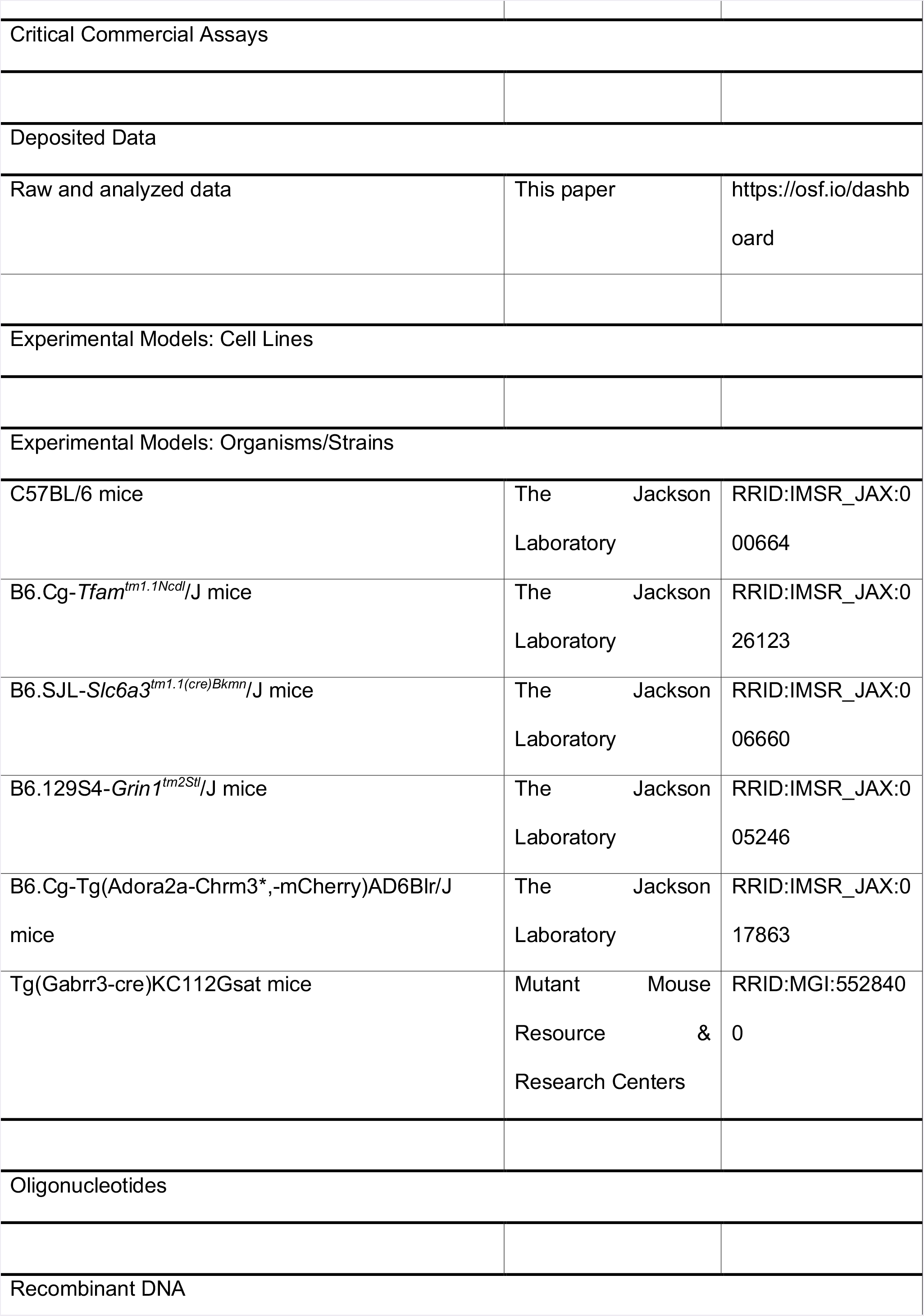
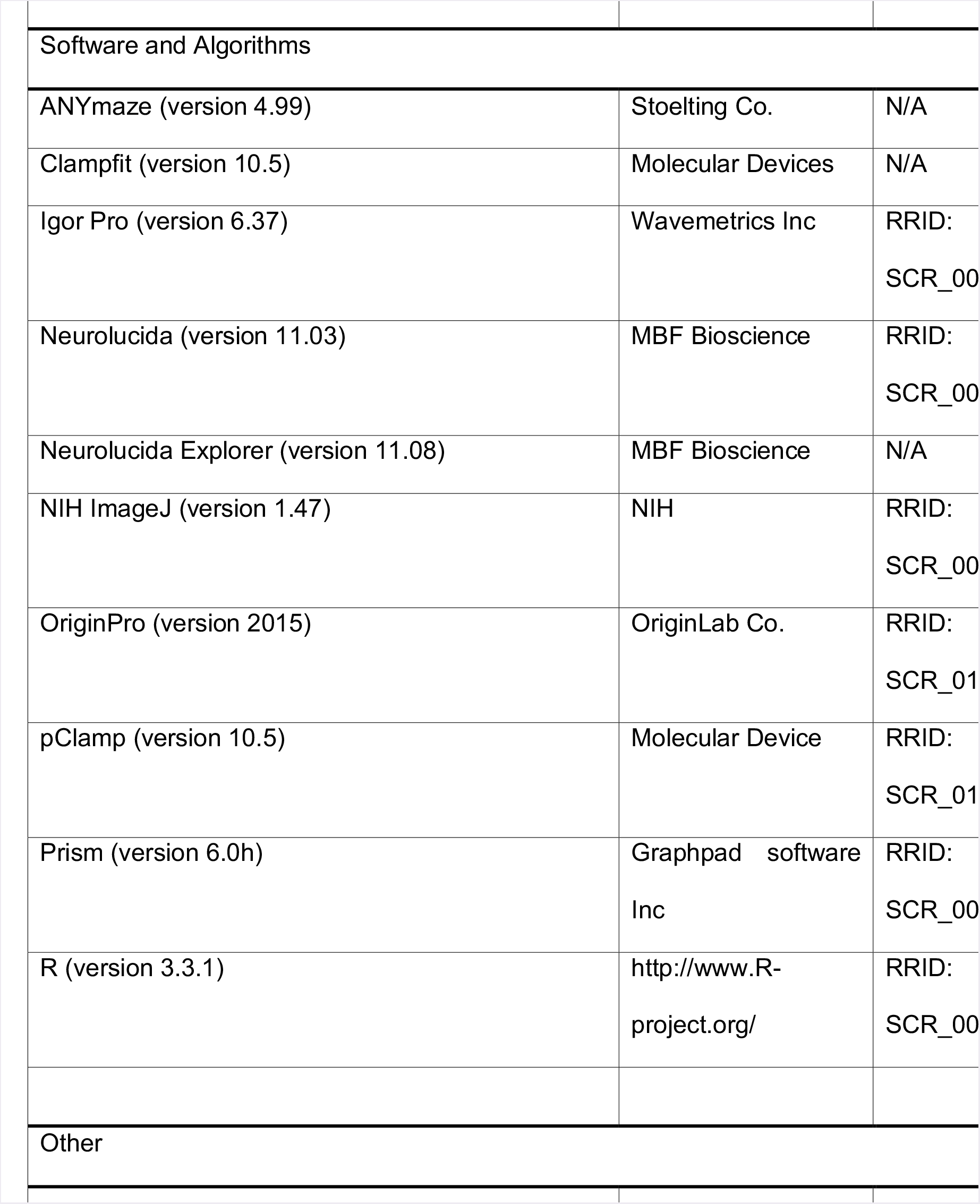
KEY RESOURCES TABLE.

## REFERENCES

Adam-Vizi, V. (2005). Production of reactive oxygen species in brain mitochondria: contribution by electron transport chain and non-electron transport chain sources. Antioxid Redox Signal 7, 1140–1149.

Albin, R.L., Young, A.B., and Penney, J.B. (1989). The functional anatomy of basal ganglia disorders. Trends NeuROSci 12, 366–375.

Atherton, J.F., and Bevan, M.D. (2005). Ionic mechanisms underlying autonomous action potential generation in the somata and dendrites of GABAergic substantia nigra pars reticulata neurons in vitro. J NeuROSci 25, 8272–8281.

Atherton, J.F., McIver, E.L., Mullen, M.R., Wokosin, D.L., Surmeier, D.J., and Bevan, M.D. (2016). Early dysfunction and progressive degeneration of the subthalamic nucleus in mouse models of Huntington’s disease. Elife 5, e21616.

Atherton, J.F., Wokosin, D.L., Ramanathan, S., and Bevan, M.D. (2008). Autonomous initiation and propagation of action potentials in neurons of the subthalamic nucleus. J Physiol 586, 5679–5700.

Baufreton, J., Atherton, J.F., Surmeier, D.J., and Bevan, M.D. (2005). Enhancement of excitatory synaptic integration by GABAergic inhibition in the subthalamic nucleus. J NeuROSci 25, 8505–8517.

Benabid, A.L., Chabardes, S., Mitrofanis, J., and Pollak, P. (2009). Deep brain stimulation of the subthalamic nucleus for the treatment of Parkinson’s disease. Lancet Neurol 8, 67–81.

Bergman, H., Wichmann, T., and DeLong, M.R. (1990). Reversal of experimental parkinsonism by lesions of the subthalamic nucleus. Science 249, 1436–1438.

Bevan, M.D., Magill, P.J., Hallworth, N.E., Bolam, J.P., and Wilson, C.J. (2002). Regulation of the timing and pattern of action potential generation in rat subthalamic neurons in vitro by GABA-A IPSPs. J Neurophysiol 87, 1348–1362.

Bevan, M.D., and Wilson, C.J. (1999). Mechanisms underlying spontaneous oscillation and rhythmic firing in rat subthalamic neurons. J NeuROSci 19, 7617–7628.

Bouabid, S., and Zhou, F.M. (2018). cAMP-producing chemogenetic activation of indirect pathway striatal projection neurons and the downstream effects on the globus pallidus and subthalamic nucleus in freely moving mice. J Neurochem 145(6): 436–448.

Chan, C.S., Shigemoto, R., Mercer, J.N., and Surmeier, D.J. (2004). HCN2 and HCN1 channels govern the regularity of autonomous pacemaking and synaptic resetting in globus pallidus neurons. J NeuROSci 24, 9921–9932.

Chu, H.Y., Atherton, J.F., Wokosin, D., Surmeier, D.J., and Bevanl, M.D. (2015). HeteROSynaptic regulation of external globus pallidus inputs to the subthalamic nucleus by the motor cortex. Neuron 85, 364–376.

Chu, H.Y., McIver, E.L., Kovaleski, R.F., Atherton, J.F., and Bevan, M.D. (2017). Loss of Hyperdirect Pathway Cortico-Subthalamic Inputs Following Degeneration of Midbrain Dopamine Neurons. Neuron 95, 1306–1318 e1305.

Cragg, S.J., Baufreton, J., Xue, Y., Bolam, J.P., and Bevan, M.D. (2004). Synaptic release of dopamine in the subthalamic nucleus. Eur J NeuROSci 20, 1788–1802.

Day, M., Wang, Z., Ding, J., An, X., Ingham, C.A., Shering, A.F., Wokosin, D., Ilijic, E., Sun, Z., Sampson, A.R., et al. (2006). Selective elimination of glutamatergic synapses on striatopallidal neurons in Parkinson disease models. Nat NeuROSci 9, 251–259.

Do, M.T., and Bean, B.P. (2003). Subthreshold sodium currents and pacemaking of subthalamic neurons: modulation by slow inactivation. Neuron 39, 109–120.

Dugan, L.L., Sensi, S.L., Canzoniero, L.M., Handran, S.D., Rothman, S.M., Lin, T.S., Goldberg, M.P., and Choi,D.W. (1995). Mitochondrial production of reactive oxygen species in cortical neurons following exposure to N-methyl-D-aspartate. J NeuROSci 15, 6377–6388.

Ekstrand, M.I., and Galter, D. (2009). The MitoPark Mouse - an animal model of Parkinson’s disease with impaired respiratory chain function in dopamine neurons. Parkinsonism Relat Disord 15 Suppl 3, S185–188.

Ekstrand, M.I., Terzioglu, M., Galter, D., Zhu, S., Hofstetter, C., Lindqvist, E., Thams, S., Bergstrand, A., Hansson, F.S., Trifunovic, A., et al. (2007). Progressive parkinsonism in mice with respiratory-chain-deficient dopamine neurons. Proc Natl Acad Sci U S A 104, 1325–1330.

Eusebio, A., Cagnan, H., and Brown, P. (2012). Does suppression of oscillatory synchronisation mediate some of the therapeutic effects of DBS in patients with Parkinson’s disease? Front Integr NeuROSci 6, 47.

Fan, K.Y., Baufreton, J., Surmeier, D.J., Chan, C.S., and Bevan, M.D. (2012). Proliferation of external globus pallidus-subthalamic nucleus synapses following degeneration of midbrain dopamine neurons. J NeuROSci 32, 13718–13728.

Farrell, M.S., Pei, Y., Wan, Y., Yadav, P.N., Daigle, T.L., Urban, D.J., Lee, H.M., Sciaky, N., Simmons, A., Nonneman, R.J., et al. (2013). A Galphas DREADD mouse for selective modulation of cAMP production in striatopallidal neurons. Neuropsychopharmacology 38, 854–862.

Fieblinger, T., Graves, S.M., Sebel, L.E., Alcacer, C., Plotkin, J.L., Gertler, T.S., Chan, C.S., Heiman, M., Greengard, P., Cenci, M.A., and Surmeier, D.J. (2014). Cell type-specific plasticity of striatal projection neurons in parkinsonism and L-DOPA-induced dyskinesia. Nat Commun 5, 5316.

Gittis, A.H., Hang, G.B., LaDow, E.S., Shoenfeld, L.R., Atallah, B.V., Finkbeiner, S., and Kreitzer, A.C. (2011). Rapid target-specific remodeling of fast-spiking inhibitory circuits after loss of dopamine. Neuron 71, 858–868.

Gradinaru, V., Mogri, M., Thompson, K.R., Henderson, J.M., and Deisseroth, K. (2009). Optical deconstruction of parkinsonian neural circuitry. Science 324, 354–359.

Hallworth, N.E., Wilson, C.J., and Bevan, M.D. (2003). Apamin-sensitive small conductance calcium-activated potassium channels, through their selective coupling to voltage-gated calcium channels, are critical determinants of the precision, pace, and pattern of action potential generation in rat subthalamic nucleus neurons in vitro. J NeuROSci 23, 7525–7542.

Hanson, G.T., Aggeler, R., Oglesbee, D., Cannon, M., Capaldi, R.A., Tsien, R.Y., and Remington, S.J. (2004). Investigating mitochondrial redox potential with redox-sensitive green fluorescent protein indicators. J Biol Chem 279, 13044–13053.

Hashimoto, T., Elder, C.M., Okun, M.S., Patrick, S.K., and Vitek, J.L. (2003). Stimulation of the subthalamic nucleus changes the firing pattern of pallidal neurons. J NeuROSci 23, 1916–1923.

Holm, S. (1979). A simple sequentially rejective multiple test procedure. Scandinavian Journal of Statistics 6, 65-70.

Ichinari, K., Kakei, M., Matsuoka, T., Nakashima, H., and Tanaka, H. (1996). Direct activation of the ATP-sensitive potassium channel by oxygen free radicals in guinea-pig ventricular cells: its potentiation by MgADP. J Mol Cell Cardiol 28, 1867–1877.

Jenkinson, N., and Brown, P. (2011). New insights into the relationship between dopamine, beta oscillations and motor function. Trends NeuROSci 34, 611–618.

Kawano, T., Zoga, V., Kimura, M., Liang, M.Y., Wu, H.E., Gemes, G., McCallum, J.B., Kwok, W.M., Hogan, Q.H., and Sarantopoulos, C.D. (2009). Nitric oxide activates ATP-sensitive potassium channels in mammalian sensory neurons: action by direct S-nitROSylation. Mol Pain 5, 12.

Krashes, M.J., Koda, S., Ye, C., Rogan, S.C., Adams, A.C., Cusher, D.S., Maratos-Flier, E., Roth, B.L., and Lowell, B.B. (2011). Rapid, reversible activation of AgRP neurons drives feeding behavior in mice. J Clin Invest 121, 1424–1428.

Kravitz, A.V., Freeze, B.S., Parker, P.R., Kay, K., Thwin, M.T., Deisseroth, K., and Kreitzer, A.C. (2010). Regulation of parkinsonian motor behaviours by optogenetic control of basal ganglia circuitry. Nature 466, 622-626.

Lee, C.R., Patel, J.C., O’Neill, B., and Rice, M.E. (2015). Inhibitory and excitatory neuromodulation by hydrogen peroxide: translating energetics to information. J Physiol 593,3431–3446.

Lemos, J.C., Friend, D.M., Kaplan, A.R., Shin, J.H., Rubinstein, M., Kravitz, A.V., and Alvarez, V.A. (2016). Enhanced GABA T ransmission Drives Bradykinesia Following Loss of Dopamine D2 Receptor Signaling. Neuron 90,824–838.

Levy, R., Lang, A.E., Dostrovsky, J.O., Pahapill, P., Romas, J., Saint-Cyr, J., Hutchison, W.D., and Lozano, A.M. (2001). Lidocaine and muscimol microinjections in subthalamic nucleus reverse Parkinsonian symptoms. Brain 124,2105–2118.

Loucif, A.J., Woodhall, G.L., Sehirli, U.S., and Stanford, I.M. (2008). Depolarisation and suppression of burst firing activity in the mouse subthalamic nucleus by dopamine D1/D5 receptor activation of a cyclic-nucleotide gated non-specific cation conductance. Neuropharmacology 55,94–105.

Magill, P.J., Bolam, J.P., and Bevan, M.D. (2001). Dopamine regulates the impact of the cerebral cortex on the subthalamic nucleus-globus pallidus network. NeuROScience 106,313–330.

Mallet, N., Ballion, B., Le Moine, C., and Gonon, F. (2006). Cortical inputs and GABA interneurons imbalance projection neurons in the striatum of parkinsonian rats. J NeuROSci 26,3875–3884.

Mallet, N., Pogosyan, A., Marton, L.F., Bolam, J.P., Brown, P., and Magill, P.J. (2008a). Parkinsonian beta oscillations in the external globus pallidus and their relationship with subthalamic nucleus activity. J NeuROSci 28,14245–14258.

Mallet, N., Pogosyan, A., Sharott, A., Csicsvari, J., Bolam, J.P., Brown, P., and Magill, P.J. (2008b). Disrupted dopamine transmission and the emergence of exaggerated beta oscillations in subthalamic nucleus and cerebral cortex. J NeuROSci 28,4795–4806.

Mathai, A., Ma, Y., Pare, J.F., Villalba, R.M., Wichmann, T., and Smith, Y. (2015). Reduced cortical innervation of the subthalamic nucleus in MPTP-treated parkinsonian monkeys. Brain 138,946–962.

Morgan, J.I., Cohen, D.R., Hempstead, J.L., and Curran, T. (1987). Mapping patterns of c-fos expression in the central nervous system after seizure. Science 237,192–197.

Mourre, C., Manrique, C., Camon, J., Aidi-Knani, S., Deltheil, T., Turle-Lorenzo, N., Guiraudie-Capraz, G., and Amalric, M. (2017). Changes in SK channel expression in the basal ganglia after partial nigROStriatal dopamine lesions in rats: Functional consequences. Neuropharmacology 113,519–532.

Nambu, A., Tokuno, H., and Takada, M. (2002). Functional significance of the cortico-subthalamo-pallidal ‘hyperdirect’ pathway. NeuROSci Res 43,111–117.

Nichols, C.G. (2006). K_ATP_ channels as molecular sensors of cellular metabolism. Nature 440,470–476.

Parker, J.G., Marshall, J.D., Ahanonu, B., Wu, Y.W., Kim, T.H., Grewe, B.F., Zhang, Y., Li, J.Z., Ding, J.B., Ehlers, M.D., and Schnitzer, M.J. (2018). Diametric neural ensemble dynamics in parkinsonian and dyskinetic states. Nature 557,177–182.

Rajakumar, N., Elisevich, K., and Flumerfelt, B.A. (1994). Parvalbumin-containing GABAergic neurons in the basal ganglia output system of the rat. J Comp Neurol 350,324–336.

Ramanathan, S., Tkatch, T., Atherton, J.F., Wilson, C.J., and Bevan, M.D. (2008). D2-like dopamine receptors modulate SKCa channel function in subthalamic nucleus neurons through inhibition of Cav2.2 channels. J Neurophysiol 99,442–459.

Roth, B.L. (2016). DREADDs for NeuROScientists. Neuron 89,683–694.

Ryan, M.B., Bair-Marshall, C., and Nelson, A.B. (2018). Aberrant striatal activity in Parkinsonism and levodopa-induced dyskinesia. Cell Rep 23,3438–3446.

Sanchez-Padilla, J., Guzman, J.N., Ilijic, E., Kondapalli, J., Galtieri, D.J., Yang, B., Schieber, S., Oertel, W., Wokosin, D., Schumacker, P.T., and Surmeier, D.J. (2014). Mitochondrial oxidant stress in locus coeruleus is regulated by activity and nitric oxide synthase. Nat NeuROSci 17,832–840.

Sanders, T.H., Clements, M.A., and Wichmann, T. (2013). Parkinsonism-related features of neuronal discharge in primates. J Neurophysiol 110,720–731.

Sanders, T.H., and Jaeger, D. (2016). Optogenetic stimulation of cortico-subthalamic projections is sufficient to ameliorate bradykinesia in 6-OHDA lesioned mice. Neurobiol Dis 95,225–237.

Schallert, T., Fleming, S.M., Leasure, J.L., Tillerson, J.L., and Bland, S.T. (2000). CNS plasticity and assessment of forelimb sensorimotor outcome in unilateral rat models of stroke, cortical ablation, parkinsonism and spinal cord injury. Neuropharmacology 39,777–787.

Sharott, A., Gulberti, A., Zittel, S., Tudor Jones, A.A., Fickel, U., Munchau, A., Koppen, J.A., Gerloff, C., Westphal, M., Buhmann, C. et al. (2014). Activity parameters of subthalamic nucleus neurons selectively predict motor symptom severity in Parkinson’s disease. J NeuROSci 34,6273–6285.

Sharott, A., Vinciati, F., Nakamura, K.C., and Magill, P.J. (2017). A Population of Indirect Pathway Striatal Projection Neurons Is Selectively Entrained to Parkinsonian Beta Oscillations. J NeuROSci 37,9977–9998.

Shen, K.Z., and Johnson, S.W. (2010). Ca2+ influx through NMDA-gated channels activates ATP-sensitive K+ currents through a nitric oxide-cGMP pathway in subthalamic neurons. J NeuROSci 30,1882–1893.

Shen, K.Z., and Johnson, S.W. (2012). Chronic dopamine depletion augments the functional expression of K-ATP channels in the rat subthalamic nucleus. NeuROSci Lett 531,104–108.

Shen, K.Z., and Johnson, S.W. (2013). Group I mGluRs evoke K-ATP current by intracellular Ca2+ mobilization in rat subthalamus neurons. J Pharmacol Exp Ther 345,139–150.

Shimamoto, S.A., Ryapolova-Webb, E.S., Ostrem, J.L., Galifianakis, N.B., Miller, K.J., and Starr, P.A. (2013). Subthalamic nucleus neurons are synchronized to primary motor cortex local field potentials in Parkinson’s disease. J NeuROSci 33,7220–7233.

Stanika, R.I., Winters, C.A., Pivovarova, N.B., and Andrews, S.B. (2010). Differential NMDA receptor-dependent calcium loading and mitochondrial dysfunction in CA1 vs. CA3 hippocampal neurons. Neurobiol Dis 37, 403-411.

Tachibana, Y., Iwamuro, H., Kita, H., Takada, M., and Nambu, A. (2011). Subthalamo-pallidal interactions underlying parkinsonian neuronal oscillations in the primate basal ganglia. Eur J NeuROSci 34,1470–1484.

Tecuapetla, F., Jin, X., Lima, S.Q., and Costa, R.M. (2016). Complementary Contributions of Striatal Projection Pathways to Action Initiation and Execution. Cell 166,703–715.

Tsien, J.Z., Huerta, P.T., and Tonegawa, S. (1996). The essential role of hippocampal CA1 NMDA receptor-dependent synaptic plasticity in spatial memory. Cell 87,1327–1338.

Vila, M., Perier, C., Feger, J., Yelnik, J., Faucheux, B., Ruberg, M., Raisman-Vozari, R., Agid, Y., and Hirsch,E.C. (2000). Evolution of changes in neuronal activity in the subthalamic nucleus of rats with unilateral lesion of the substantia nigra assessed by metabolic and electrophysiological measurements. Eur J NeuROSci 12, 337-344.

Walters, J.R., Hu, D., Itoga, C.A., Parr-Brownlie, L.C., and Bergstrom, D.A. (2007). Phase relationships support a role for coordinated activity in the indirect pathway in organizing slow oscillations in basal ganglia output after loss of dopamine. NeuROScience 144,762–776.

Wang, X., and Michaelis, E.K. (2010). Selective neuronal vulnerability to oxidative stress in the brain. Front Aging NeuROSci 2, 12.

Wang, X., Zaidi, A., Pal, R., Garrett, A.S., Braceras, R., Chen, X.W., Michaelis, M.L., and Michaelis, E.K. (2009). Genomic and biochemical approaches in the discovery of mechanisms for selective neuronal vulnerability to oxidative stress. BMC NeuROSci 10, 12.

West, M.J. (1999). Stereological methods for estimating the total number of neurons and synapses: issues of precision and bias. Trends NeuROSci 22,51–61.

Wichmann, T., Bergman, H., and DeLong, M.R. (2018). Basal ganglia, movement disorders and deep brain stimulation: advances made through non-human primate research. J Neural Transm (Vienna) 125, 419–430.

Wilson, C.J. (2013). Active decorrelation in the basal ganglia. NeuROScience 250,467–482.

Wilson, C.L., Cash, D., Galley, K., Chapman, H., Lacey, M.G., and Stanford, I.M. (2006). Subthalamic nucleus neurones in slices from 1-methyl-4-phenyl-1,2,3,6-tetrahydropyridine-lesioned mice show irregular, dopamine-reversible firing pattern changes, but without synchronous activity. NeuROScience 143,565–572.

Yoon, H.H., Park, J.H., Kim, Y.H., Min, J., Hwang, E., Lee, C.J., Suh, J.K., Hwang, O., and Jeon, S.R. (2014). Optogenetic inactivation of the subthalamic nucleus improves forelimb akinesia in a rat model of Parkinson disease. NeuROSurgery 74,533–540; discussion 540–531.

Zaidel, A., Arkadir, D., Israel, Z., and Bergman, H. (2009). Akineto-rigid vs. tremor syndromes in Parkinsonism. Curr Opin Neurol 22,387–393.

Zhang, D.M., Chai, Y., Erickson, J.R., Brown, J.H., Bers, D.M., and Lin, Y.F. (2014). Intracellular signalling mechanism responsible for modulation of sarcolemmal ATP-sensitive potassium channels by nitric oxide in ventricular cardiomyocytes. J Physiol 592,971–990.

Zhu, Z., Bartol, M., Shen, K., and Johnson, S.W. (2002a). Excitatory effects of dopamine on subthalamic nucleus neurons: in vitro study of rats pretreated with 6-hydroxydopamine and levodopa. Brain Res 945,31–40.

Zhu, Z.T., Shen, K.Z., and Johnson, S.W. (2002b). Pharmacological identification of inward current evoked by dopamine in rat subthalamic neurons in vitro. Neuropharmacology 42,772–781.

Zold, C.L., Ballion, B., Riquelme, L.A., Gonon, F., and Murer, M.G. (2007). NigROStriatal lesion induces D2-modulated phase-locked activity in the basal ganglia of rats. Eur J NeuROSci 25,2131–2144.

